# Cryo-electron microscopy structure of the bovine ephemeral fever virus RNA-nucleoprotein assembly

**DOI:** 10.64898/2026.06.26.734764

**Authors:** Anastasiia Herman, Alfred A. Antson, Pavol Bardy

**Affiliations:** York Structural Biology Laboratory, Department of Chemistry, University of York, York YO10 5DD, United Kingdom; York Biomedical Research Institute, University of York, York, United Kingdom

## Abstract

Bovine ephemeral fever virus (BEFV), a member of the *Rhabdoviridae* family, is an arthropod-borne pathogen that causes acute febrile disease in cattle. The structural basis of its genome encapsidation and virion assembly remains unexplored, with the current knowledge largely limited to predictions derived from bioinformatic comparisons with other rhabdoviruses. Furthermore, the structural principles that permit the formation of variable-diameter nucleocapsids resulting in the distinctive bullet-shaped morphology of rhabdoviruses remain poorly understood. Here, we report the cryo-electron microscopy structure of the BEFV nucleoprotein (N) in complex with RNA, in the absence of other viral components. The complex predominantly forms circular decameric oligomers that we propose to act as nucleation intermediates during assembly of the bullet-shaped nucleocapsids. Direct subunit interactions are limited to a small polar surface area, with additional intersubunit links mediated by flexible N- and C-terminal loops. These interfaces generate a structurally plastic oligomeric lattice in which neighbouring N subunits can undergo substantial rigid-body rotations and positional rearrangements while preserving conserved local contacts and continuous RNA encapsidation. Such quasi-equivalent interactions provide a plausible mechanism for accommodating the progressive changes in helical diameter required for the transition from the highly curved bullet tip to the wider cylindrical trunk of rhabdovirus nucleocapsids. The assembly is stabilised by the bound RNA molecule, where nine RNA bases are accommodated by each N subunit. The RNA-binding mechanism is consistent with that of VSV, the closest BEFV homologue characterised structurally, but differs at about half of the RNA-binding residues, demonstrating the versatility of the nucleoprotein scaffold in interacting with ssRNA. Comparative analysis with other rhabdoviruses, as well as negative-sense RNA viruses with constant-diameter nucleocapsids, such as Ebola, further confirms the structural features that enable bullet-shaped versus cylindrical nucleocapsid assembly.

## Introduction

RNA viruses are the most prevalent pathogens of animals and humans. Their ability to cause large-scale outbreaks has an immense impact on global health, food production, and thus economic stability. One such virus is bovine ephemeral fever virus (BEFV), a member of the group of negative-sense RNA viruses. It infects cattle and water buffalo, causing an ephemeral fever disease, also referred to as three-day disease [1,2]. Its clinical signs include high fever, lameness, muscle weakness and limb paralysis, resulting in reduced milk production and, in severe cases, death. Ephemeral fever disease is widespread across Africa, South Asia and Australia [3], with one of the most recent outbreaks reported in February 2024 [4]. Transmission occurs through insect vectors, particularly biting midges and mosquitoes.

BEFV belongs to the order *Mononegavirales*, the family *Rhadboviridae*, and the genus *Ephemerovirus* [5]. Morphologically, BEFV is an enveloped virus with a characteristic bullet-shaped virion [1,2], a defining feature of many rhabdoviruses. The virion is formed by five structural proteins shared among *Rhabdoviruses*: the nucleoprotein (N), matrix protein (M), glycoprotein (G), phosphoprotein (P), and the large multifunctional enzyme (L), which includes an RNA-dependent RNA polymerase domain [5–7].

The nucleoprotein, hereafter referred to as N protein, plays a central role in the viral life cycle [7–9]. It interacts with the viral negative-sense genomic RNA to form a helical ribonucleoprotein complex stored in the core of the virion, stabilised by interactions with the two layers of matrix protein that link the ribonucleoprotein with the outer lipid bilayer of the virion and the surface glycoprotein [10,11]. The N protein also interacts with the P protein, which is associated with the principal replication component, the L protein that drives the synthesis of mRNA required for protein production, thus triggering production of the ‘building blocks’ for progeny virions [8,12,13]. The viral lifecycle begins with the uncoating of the virion once the trimeric receptor-binding G protein recognises the host cell. Removal of the viral membrane envelope during fusion with the host membrane allows the N protein-encapsidated (-)ssRNA genome to enter the host cytoplasm, where it becomes accessible for transcription and replication [7–10].

The N protein subunit comprises two domains, an N-terminal domain or N-lobe and a C-terminal domain or C-lobe [14–19], as seen across all studied species of the *Mononegavirales* order, including major pathogens such as Ebola (*Filovirida*e family) [20], Nipah (*Paramyxoviridae* family) [21], Measles (*Paramyxoviridae* family) [22] and Respiratory Syncytial Virus (*Pneumoviridae* family) [23]. Multiple N subunits bind to the genomic ssRNA to form a helical assembly, which is the basis of the bullet-shaped virion [24]. The genomic ssRNA resides in a cleft between the N- and C-lobes. Notably, the locations of the RNA bases, the number of nucleotides per protein subunit, and the conformation of the bound ssRNA vary among viral families.

Despite the central role of the N protein and the economic importance of BEFV, no structural characterisation of the protein-RNA complex has yet been conducted for any *Ephemerovirus* species. The current understanding of the structure can only be inferred from previous structural studies of archetypal *Rhabdoviridae* members, vesicular stomatitis virus (VSV) [16–19,25] and rabies virus (RABV) [14,26,27]. These viruses, however, are only distantly related to BEFV, with their N proteins sharing 31% and 23% sequence identity with the BEFV N protein, respectively.

Here, we report a high-resolution cryo-electron microscopy (cryo-EM) structure of the BEFV nucleoprotein in complex with ssRNA. The structure reveals an extensive network of hydrogen bonding interactions between the nucleoprotein and the sugar-phosphate backbone of RNA, defining the molecular basis of genome encapsidation. The structure also delineates intersubunit interactions that drive nucleocapsid assembly. The N- and C-terminal segments, historically and hereafter referred to as “N-terminal arm” and “C-loop”, insert into defined recognition pockets of neighbouring protomers, forming interlocking, clasp-like interactions that stabilise the helical assembly. Comparative structural analysis with N-proteins from other members of *Mononegarivales* reveals a striking evolutionary pattern: while N-RNA interactions are relatively conserved within the family, intersubunit contacts are considerably more variable and appear to be genus-specific. This divergence suggests that RNA recognition imposes strict structural constraints, whereas oligomerisation interfaces provide a platform for evolutionary adaptation, and thanks to the flexibility of the intersubunit interlocks, act as a structural determinant of the bullet-shaped virion.

## Material and methods

### Production and purification of BEFV nucleoprotein

A sequence of the BEFV N gene (GenBank NC_002526.1, 51–1378) [28] was codon-optimised for *Escherichia coli*, and synthesised into the plasmid pET28a to contain a cleavable N-terminal His-tag (GenScript). The plasmid was transformed into *E. coli* strain BL23(DE3) and cells were grown in LB liquid media containing 30 μg/ml of kanamycin at 37 ℃. When the culture reached OD_600_ = 0.6, it was cooled on ice for 30 min, and isopropyl-β-d-thiogalactopyranoside (IPTG) was added to a final concentration of 0.1 mM to induce protein production. The gene was expressed at 16 ℃ overnight. Cells were spun, and the pellets were sonicated in the presence of a protein buffer containing 20 mM Tris-HCl, pH 7.5; 300 mM NaCl. Then, disrupted cells were centrifuged at 14000 g, 10 min at 4 ℃, and the supernatant containing soluble protein was collected. The presence of the soluble protein of expected size was verified by SDS-PAGE (**Figure S1**).

For protein purification, the supernatant containing soluble BEFV nucleoprotein was applied onto a 5 ml Ni HisTrap column. The column was equilibrated using an equilibration buffer containing 20 mM TrisHCl, pH 7.5; 300 mM NaCl; 20 mM imidazole. Then, BEFV nucleoprotein was eluted using a linear gradient of an elution buffer containing an increased concentration of 500 mM imidazole. The eluted protein sample was concentrated and buffer-exchanged into a 20 mM imidazole-containing buffer using tubes with cellulose filters and 10 kDa pores (Amicon Ultra, Merck Millipore). Then, the His-tag cleavage was performed using thrombin (10 U per mg of protein) at 4 ℃ overnight. A cleaved protein sample was used to perform a His-tag pull-down using Ni-agarose beads (Amintra, Abcam) (1 ml of beads per 50 mg of protein). The protein sample free of His-tag was concentrated and exchanged into the buffer containing 20 mM TrisHCl, pH 7.5; 300 mM NaCl using the filter tubes. Further purification was done by size exclusion chromatography using a Superose 6 Increase 10/300 GL column, resulting in an elution peak of 14.8 ml, from which fractions were collected (**Figure S1**). Protein concentration was measured using NanoDrop readings at A_280nm_, and the protein was flash-frozen in liquid nitrogen and stored at -80 ℃.

### Electron microscopy sample preparation and data collection

For negative-stain EM, the BEFV nucleoprotein sample at a concentration of 0.008 mg/ml in a volume of 4 μl was applied to glow-discharged Formvar/Carbon 200 Mesh Copper grids (Agar Scientific). Grids were stained with 2% uranyl acetate and imaged using a transmission electron microscope Tecnai 12 BioTWIN G2, at 120 keV with SIS Megaview III CCD camera. For cryo-EM, four μl of BEFV nucleoprotein sample in a concentration of 0.8 mg/ml was applied on glow-discharged UltraAuFoil R1.2/1.3 with or without 0.01% of poly-l-lysine. The sample was blotted for 2 sec, -10 force at 100% humidity using Vitrobot Mark IV (ThermoFisher Scientific) and plunge-frozen in liquid ethane. Two datasets were collected using a Glacios cryogenic transmission electron microscope (ThermoFisher Scientific) equipped with a Falcon4 camera, operating at 200 keV (**Table S1**). Overnight data collection was set up using EPU software. The imaging was done at the University of York.

### Image processing

The single particle analysis of cryoEM data was performed using RELION5 [29] as shown in **Figure S2**. Two collected data sets of raw movies were imported as two different optic groups and motion corrected separately using the MotionCor2 implementation in RELION [30]. CTF estimation was performed using CTFFIND4 [31]. RELION5 implementation of Topaz [32] was used for automatic particle picking, trained on 200 manually picked particles. The obtained particle set was subjected to the reference-free 2D classification into fifty classes. The initial model was generated *de novo* with C1 symmetry applied and, after revealing decamer-like density, resampled to coincide with the C10 symmetry axis using ChimeraX [33]. This map was used as a reference for 3D refinement with C10 symmetry applied. The refined particles were subjected to several rounds of 3D classification. CTF refinement and Bayesian polishing, and a second round of CTF refinement were done using RELION5 [29]. This led to a map of 2.9 Å resolution (**Figure S3**). Local resolution analysis was performed using RELION5’s own implementation.

### Model building, refinement and interaction analysis

BEFV N decameric structure was predicted using AlphaFold3 online server [34] with predicted template modelling value of 0.66 and interface predicted template modelling value of 0.63. The N monomer was extracted and fitted into the obtained density map using the ‘fit’ command in ChimeraX [33]. RNA was predicted as a poly-uridine chain in complex with three BEFV N monomers using the AlphaFold3 online server [34], where the poly-U chain was extracted and fitted into the density map for further model building and refinement, together with the protein model.

Model building of the BEFV ribonucleoprotein complex was performed using Coot [35], and the refinement of the model was done using phenix.real_space_refine within Phenix [36], Refmac Servalcat within the CCPEM 1.7.0 suite [37] and ISOLDE 1.9 [38] (**Table S2**). Electrostatic surface potential visualisation was done using APBS-PDB2PQR server [39] and ChimeraX [33]. The analysis of protein-protein and protein-RNA interaction was done using the PDBePisa tool [40,41]. An integrated structural overlay analysis of protein-RNA interactions for selected *Mononegavirales* species was performed using PDBePisa [40,41], DALI [42] and ChimeraX [33].

### Bioinformatic, phylogenetic analysis and visualisation

A phylogenetic analysis of the nucleoprotein was performed using amino acid sequences of selected *Mononegavirales* species, aligned with ClustalW [43]. A phylogenetic tree was constructed using RAxML and the PROTGAMMA substitution model with a GTR matrix [44], followed by a bootstrap analysis using the BOOSTER web service [45] with 100 replicates. A multiple sequence alignment (MSA) was performed using the ESPript 3.0 online tool [46,47]. The phylogenetic tree was visualised using MEGA12 [48]. The figures used in this article were created using ChimeraX [33] and BioRender (https://BioRender.com).

## Results

### Overall structure of the nucleoprotein-RNA complex

The 52kDa BEFV nucleoprotein was overexpressed in *E.coli*. Size-exclusion chromatography elution peak of the purified protein indicated the molecular weight of ∼500 kDa (**Figure S1**), consistent with a 9-10 subunit assembly. The negative stain TEM showed multimeric ring-like structures (**Figure S1**). The 260/280nm absorption ratio of above 1 suggested that the purified protein complex retained associated nucleic acid (**Figure S1**). Single-particle analysis (SPA), based on the first cryo-EM data set (**Table S1**), showed a predominant presence of decameric rings, but with significant preferential orientation of the particles, with few side views (**Figure 1A, Figure S2**). The addition of a low concentration of poly-L-lysine improved particle distribution, and a second dataset displayed markedly improved orientation diversity as seen in the 2D class averages (**Figure 1A, 1B**). Both data sets (**Table S1, Figure S2**) were merged and used for further 3D reconstruction. The 2D classification revealed some heterogeneity among protein assemblies, with ∼91% majority of particles forming decameric rings, while 11-mer and 12-mer oligomers each accounted for only ∼4.5% of particles, as estimated from the top views (**Figure S3**). Further reconstruction was based on the prevalent decameric top-view and all side-view classes. An initial model, generated *de novo* in C1 (**Table S1, Figure S2**), showed no visible deviations from the decameric structure and hence, C10 symmetry was imposed for subsequent processing of the merged data set.

**Figure 1.**
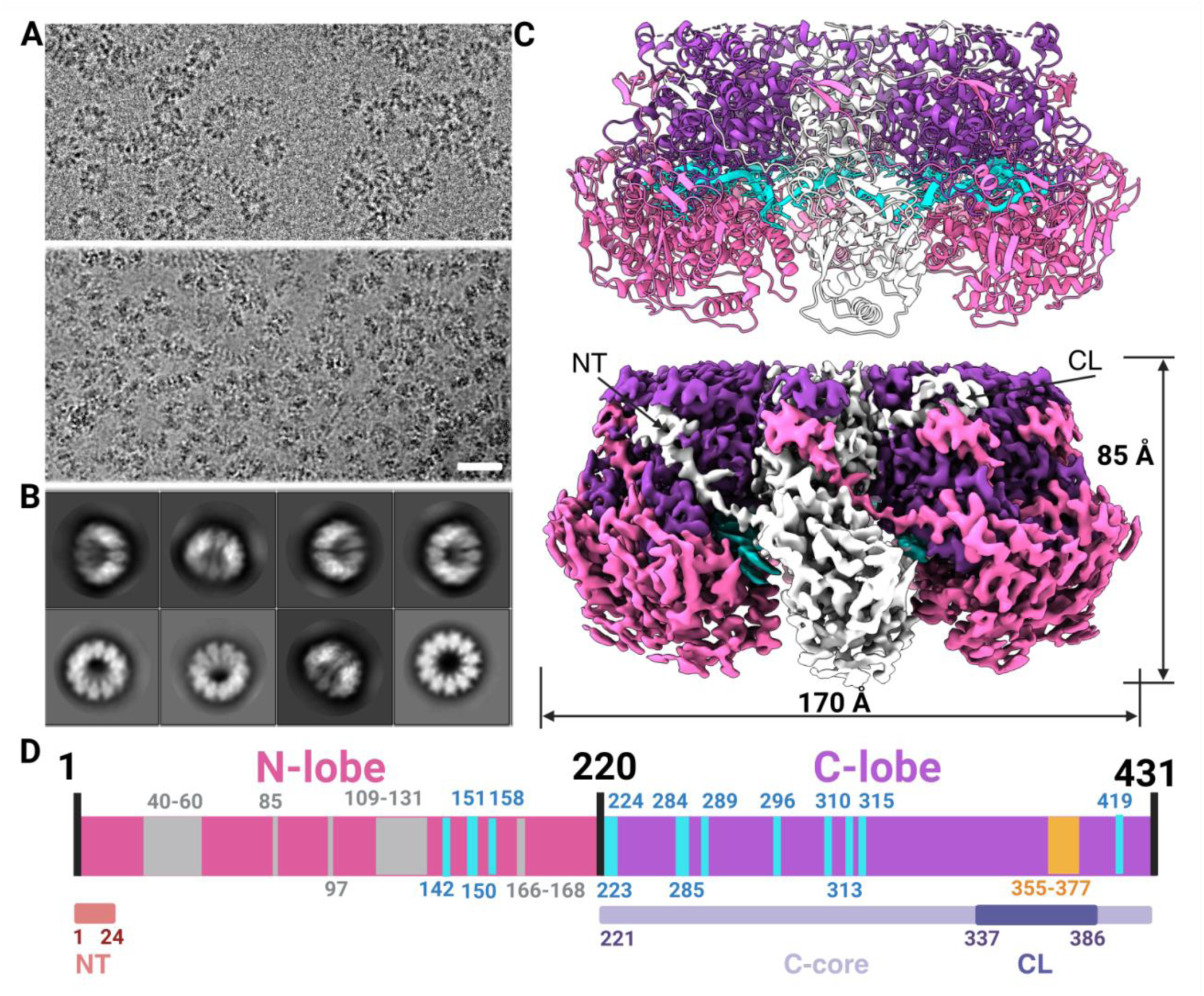
Overall structure of the BEFV nucleoprotein. (A) Cryo-electron micrographs of BEFV nucleoprotein from dataset 1, showing preferential orientation (top), and from dataset 2, collected on poly-lysine-coated grids, which improved particle distribution (bottom). Scale bar, 50 nm. (B) Selected 2D class averages. (C) Top: Model of the decameric nucleoprotein assembly with ssRNA in cyan, C-lobe in purple and N-lobe in pink. One subunit is shown in white. Bottom: cryo-EM density map of the complex, coloured as above. N-terminal arm (NT) and the C-loop (CL) of the central subunit (white) are labelled. (D) Schematic of nucleoprotein subunit with N- and C-lobe boundary residues labelled. Regions with diffused density are in grey, and the unmodelled segment is in yellow. Residues interacting with RNA, identified by PDBePISA [40,41], are numbered in light blue.

Single particle 3D reconstruction resulted in a 2.9 Å resolution density map (**Figure S2, Figure S3**), revealing a decameric circular assembly with the overall height of 85 Å, outer diameter of 170 Å and inner diameters of 130 Å and 48 Å for the widest and most constricted parts, respectively (**Figure 1C**). In common with nucleoproteins of other studied *Rhabdoviridae,* each subunit can be subdivided into N and C-lobe domains [14,16–21]. The N-lobe consists of residues 1-220 comprising α-helices 1-9, 3_10_ helices 1-4 and antiparallel and parallel β-strands 1-6. The C-lobe includes residues 221-431 containing α-helices 10-20 and 3_10_ helices 5-6 (**Data S1**). Cryo-EM density for residues 40-80, 100-130 and 160-170 of the N-lobe is relatively diffused, indicating flexibility of these regions (**Figure 1D, Figure S4, Figure S5**). Model building for these segments was guided by the AlphaFold3-predicted model. The C-lobe has only one region with diffused density, which is least well-defined, corresponding to an extended exposed loop, residues 355-377. This loop was unmodelled due to the lack of defined density (**Figure 1D**). Hereafter, we will refer to the C-lobe region spanning amino acids 338-386 as the C-loop, adopting the terminology commonly used for description of this segment of the nucleoprotein in other members of the *Rhabdoviridae* [14,16–21].

### Protein-RNA interactions

The cryo-EM density map revealed ssRNA bound along a groove at the internal surface area of the oligomeric protein ring. Local resolution within the RNA-binding groove reached 2.7 Å (**Figure S5**), featuring clear density corresponding to ssRNA, which co-purified with the recombinant protein. The ssRNA was modelled as a poly-U chain (**Table S3**) to account for non-specific protein-RNA interactions. Consistent with other members of the *Rhabdoviridae*, each N protein subunit binds nine RNA nucleotides (**Figure 2A**) inside the positively charged cleft (**Figure S6)**, formed between the C and N-lobes (**Figure 2B**). The 9 RNA nucleotides adopt the characteristic ‘3-out, 1-in, 1-out, 2-in, 2-out’ conformation previously observed in VSV and RABV nucleoproteins.

**Figure 2.**
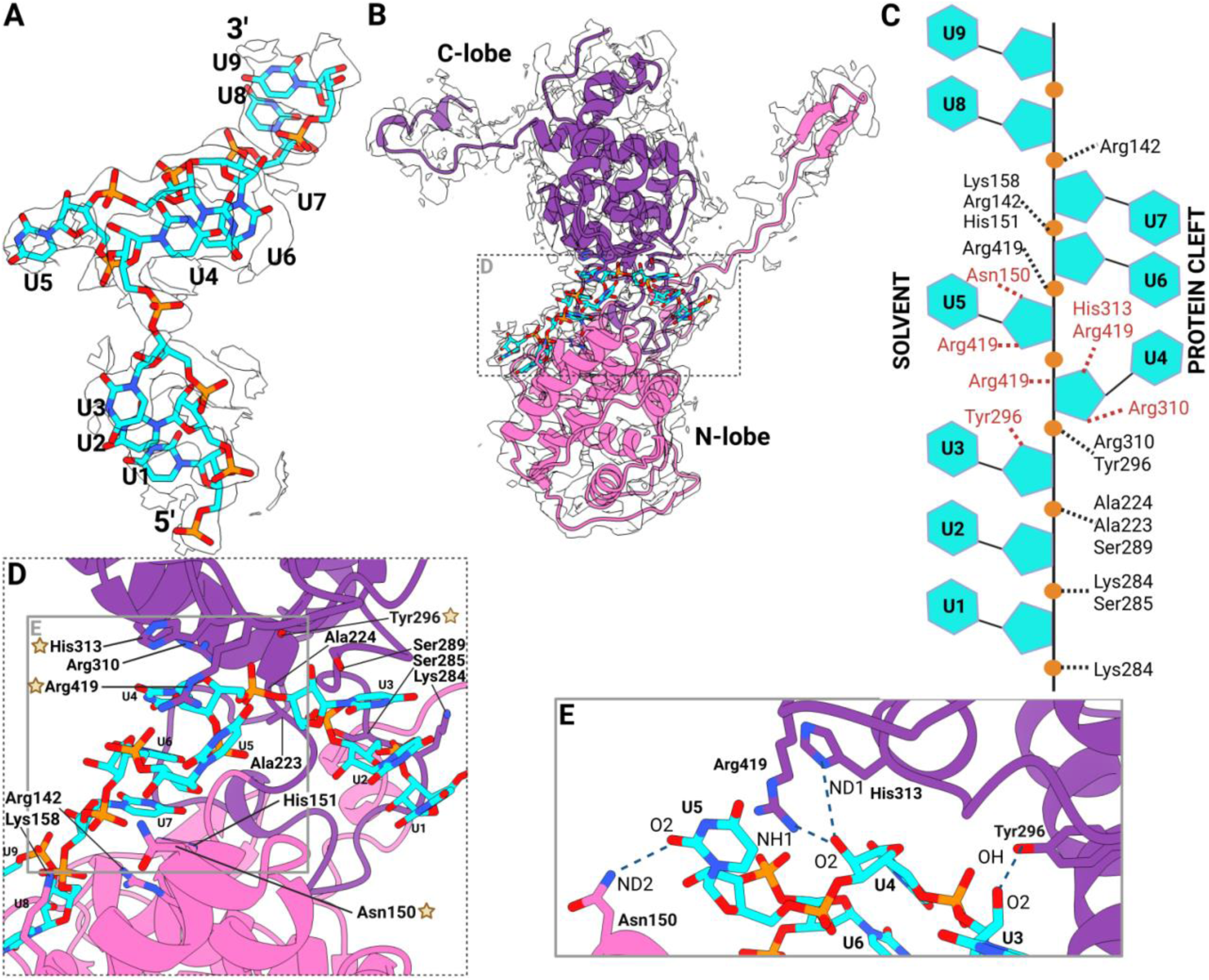
Protein-RNA interactions. (A) Cryo-EM density corresponding to bound RNA (white) with fitted poly-U model (RNA C, O, N and P atoms in cyan, red, blue and orange, respectively). (B) Cryo-EM density map (white) for a nucleocapsid protomer with corresponding models of protein (ribbon) and RNA (sticks). (C) Schematic representation of RNA chain with protein-RNA hydrogen bonding and ionic interactions. Residues interacting with the RNA phosphate and sugar moieties are in black and red, respectively. (D) Close-up view of the RNA-binding cleft, with side chains of interacting amino acids shown in sticks. Residues interacting with the 2’-OH groups of the RNA are marked with yellow stars. (E) Hydrogen bonding interactions involving 2’-OH groups of RNA, viewed at 45° around the z-axis compared to panel (D).

RNA binding is mediated by an extensive network of hydrogen bonding and ionic interactions involving 14 amino acids from both lobes of each BEFV nucleoprotein subunit (**Figure 2D, 2E, Table S4**). Protein-RNA interactions are dominated by electrostatic contacts with the sugar-phosphate backbone, rather than specific interactions with the bases. This indicates that RNA binding is driven primarily by charge complementarity. In particular, positively charged residues including Arg142, Lys158, Arg310 and Arg313, form multiple salt bridges with RNA phosphates, generating an extended electropositive surface within the binding cleft. Arg142 is particularly important, forming three salt bridges with phosphates of nucleotides U7 and U8 (**Figure 2C**, **Table S4**). Additional stabilising interactions are formed with several polar residues, including His151, Ser285, Ser289, Tyr296 and His419, which form hydrogen bonds with the RNA (**Figure 2C, 2D, Table S4**). Several interactions involve the 2′-OH groups of ribose sugars, particularly for nucleotides U3, U4, and U5, conferring specificity for RNA rather than DNA (**Figure 2C**, **Table S4**). Among these nucleotides, U4 forms the most extensive interaction network, including contacts with 2՛-OH, 3՛-OH and 4՛-OH hydroxyls, as well as stacking interactions with bases of nucleotides 6 and 7 (**Figure 2C, Table S4**).

The prevalence of positively charged RNA-binding residues is consistent with observations in other *Mononegavirales* species [20,25,26], where negatively charged residues are generally absent from the RNA-binding cleft. This finding emphasises the importance of ionic interactions in non-sequence-specific encapsidation of the ssRNA genome. Despite significant sequence differences, the overall mode of RNA binding and its trajectory are conserved across structurally characterised rhabdoviruses. The reported structures of N-RNA complexes of VSV [25] and RABV [26] have essentially identical protein-RNA architectures, and, like BEFV, bind nine nucleotides per N subunit (**Figures 3A, 3C, 3D**). This contrasts with more distant members of *Mononegavirales*, including filoviruses (**Figure 3E**), paramyxoviruses and pneumoviruses, where six nucleotides per subunit are predominant [21–23]. Thus, the nine-nucleotide register and RNA trajectory observed for BEFV appear to be the defining structural feature of *Rhabdoviridae*.

**Figure 3.**
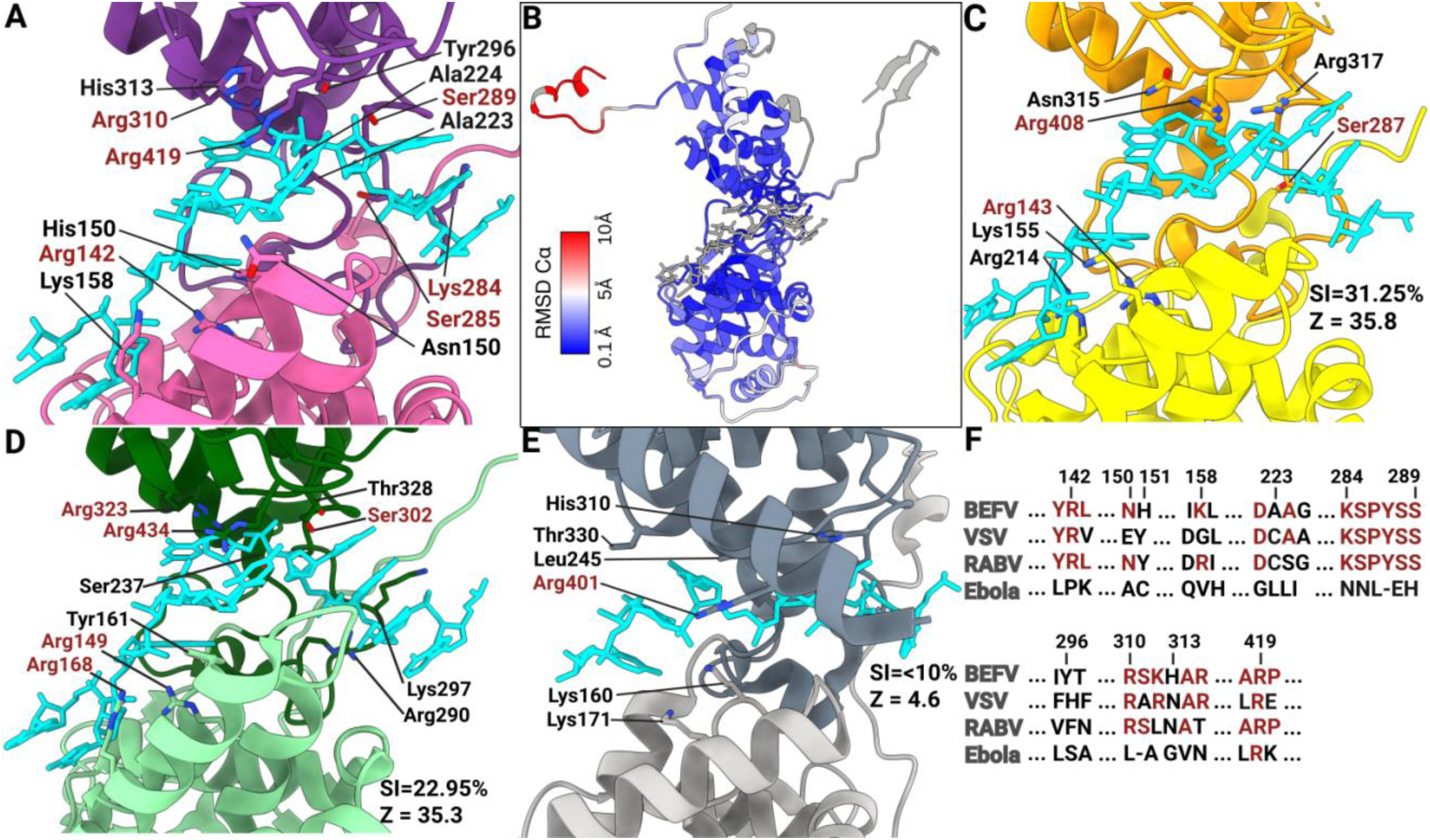
**Comparison of RNA-binding in nucleoproteins from different members of the *Mononegavirales.*** Proteins are shown as ribbons with side chains of RNA-binding residues in sticks. RNA is in cyan. (A) BEFV N-RNA complex with conserved residues highlighted in red. (B) BEFV nucleoprotein coloured based on the RMS deviation from Cα atoms of VSV N. Non-homologous protein regions and the ssRNA are in grey. (C-E) VSV N-RNA complex (C, pdb:7UWS), RABV N-RNA complex (D, pdb:8FFR) and Ebola N-RNA complex (E, pdb:5Z9W) with conserved residues highlighted in red. Sequence identity (SI) and Z-scores calculated using DALI [42] are shown at the bottom right of each panel. (F) Structure-based sequence alignment of the RNA-binding residues of BEFV, VSV, RABV and Ebola. Residue numbering corresponds to BEFV N with conserved residues marked in red. Three dots reflect sequence segments not shown on the figure, dashes depict the absence of corresponding residues.

To analyse the conservation of the RNA interaction groove, a multiple sequence alignment was generated from 13 *Ephemerovirus* species, along with VSV and RABV as representatives of the wider *Rhabdoviridae* family, and 4 additional members of the wider *Mononegavirales* order (**Data S1**). Additionally, structure-based comparison of RNA-binding residues of nucleoproteins from four species with available structures, including VSV, RABV and Ebola, was performed using DALI [42]. A separate sequence conservation analysis was also performed using all twenty sequenced *Ephemerovirus* species. These analyses showed that the RNA-binding groove is the most highly conserved region of the nucleoprotein (**Figure S7**).

Interestingly, only a single RNA-binding residue, Arg419, is conserved across all analysed *Mononegavirales* species (**Figure 3, Data S1**). In all structurally characterised homologues, including VSV, RABV, and Ebola virus, the equivalent arginine forms multiple hydrogen bonds and salt bridges with RNA (**Figure 3**), indicating its conserved functional importance for RNA binding. In consonance, Arg419 makes the highest free energy contribution (ΔG -1.35 kcal/mol) among BEFV N residues involved in RNA binding.

Within the *Rhabdoviridae* family, five additional RNA-binding residues are conserved: Arg142, Lys284, Ser285, Ser289, and Arg310 (**Figure 3**). Arg142 corresponds to Arg143 in VSV and Arg149 in RABV, retaining its ionic interaction with RNA phosphate. Likewise, Lys284 corresponds to Lys297 in RABV, where it also forms the salt bridge with RNA (**Figure 3A, 3D, 3F**). VSV contains structurally conserved Lys286, as seen in the crystal structure (pdb: 2gic), although in the deposited cryo-EM structure (pdb: 7uws), the side chain of this residue is oriented away from RNA. This discrepancy may reflect differences between the recombinantly produced form (pdb: 2gic) and mature virions (pdb: 7uws), or alternative conformations of this residue. Arg310 retains its ionic interaction with RNA in RABV but not in VSV (**Figure 3**). Considering residues involved in hydrogen bonding with RNA, there is more variation among *Rhabdoviridae* species. In particular, interactions mediated by Ser285 and Ser289 are differentially retained in RABV and VSV (**Figure 3**). Other RNA-binding residues of BEFV exhibit varying degrees of conservation (**Data S1**). However, the overall mechanism of RNA binding is preserved, with binding relying primarily on hydrogen bonds and ionic interactions with the sugar-phosphate backbone, with no structural evidence of nucleotide-specific recognition. Interestingly, BEFV N RNA-binding residues are more structurally conserved with RABV than with VSV, despite a closer phylogenetic relationship with VSV (**Data S1**). Notwithstanding, the overall distribution of polar and hydrophobic surfaces within the RNA-binding groove of N is similar across *Rhabdoviridae* members (**Figure S8**).

Together, these observations show that the high similarity in RNA binding observed among *Rhabdoviridae* members is largely achieved through conservation of the overall architecture and electrostatic complementarity, with a strict conservation of only a few individual residues. Such a combination of structural conservation with significant sequence variation enables different rhabdoviruses to maintain efficient genome encapsidation while allowing significant evolutionary divergence.

### Protein-protein interaction between nucleoprotein subunits

The nucleoprotein assembly is stabilised by subunit-subunit interactions with the total buried surface area of 2,140 Å^2^ per protomer. There are three dominant regions within this interface: the C-core region of the C-lobe, the C-loop (CL) and the N-terminal arm (NT) (**Figure 4)**. Together, these regions form an interconnected network, where the C-core of a central subunit is linked to C-cores of adjacent subunits and to both the NT of the *n-1* subunit, as well as CL of the *n+1* subunit (**Figure 4**). Another interaction interface forms between the NT region of the *n-1* subunit and the CL of the *n+1* subunit. These interactions include a salt bridge between the Lys7 and Glu355 side chains, as well as six hydrogen bonds (**Table S5, Figure 4B**).

**Figure 4.**
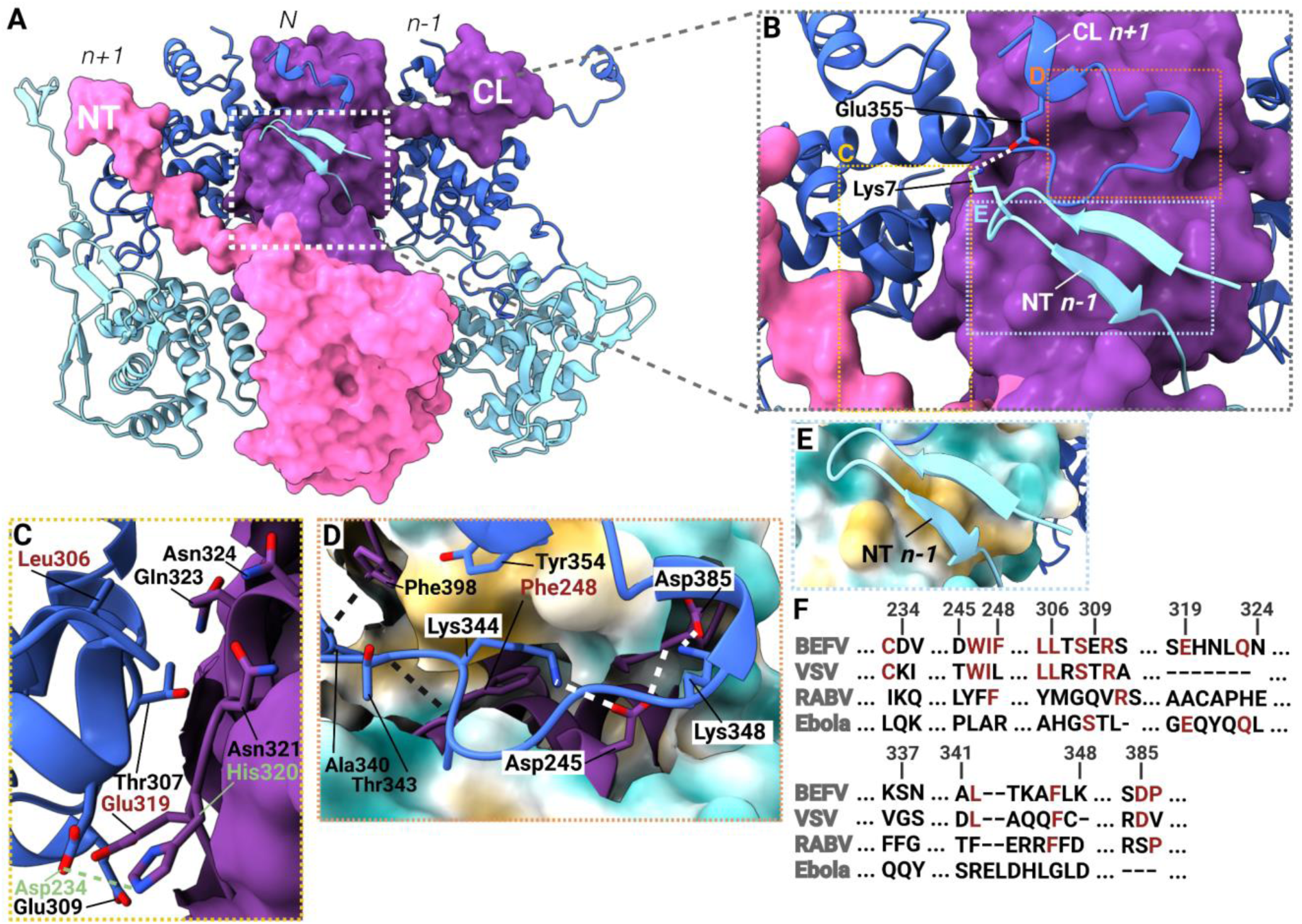
Subunit interactions. (A) Three adjacent subunits, where the central is shown as a surface with the N-lobe in pink and the C-lobe in purple. Neighbouring (*n+1* and *n-1*) subunits are shown as ribbons with N-lobes in light blue and C-lobes in dark blue. (B) Interactions between *n+1* and *n-1* subunits, with side chains of residues forming salt bridges (white dash) shown as sticks. (C) Interactions between adjacent C-lobes with residues forming ionic interactions labelled in green. Conserved and non-conserved residues forming H-bonds are labelled in red and black, respectively. (D) Interactions between the C-core of the central subunit (molecular surface) and the CL of the *n+1* subunit (ribbon). Hydrogen bond-forming residues are shown and labelled in black, residues forming ionic interactions (white dashed lines) are highlighted in white rectangles. The molecular surface colour is changing from blue (hydrophilic) to yellow (hydrophobic). (E) Interactions between the C-core of *N* subunit (molecular surface) with the N-terminal arm of the *n-1* subunit (ribbon). (F) Structure-based sequence alignment of BEFV, VSV, RABV and Ebola nucleoproteins generated by DALI [42], for residue segments shown in panels B-D. Conserved residues are in red, dashes indicate gaps, and dots denote regions absent from the figure.

Contacts between the C-core of the *N* subunit and the NT and CL regions of adjacent subunits result in an extensive interaction surface (**Figure 4B, 4D, 4E**), likely critical for nucleoprotein oligomer self-assembly. These interfaces are dominated by hydrophobic interactions formed by C-core residues Ile226, Leu226, Leu229, Ile240, Ile247, Leu344, Met260, Ile266 with the NT, and C-core residues Phe248, Met327, Leu393, Phe398, Met387, Ile391, Leu400 with the CL (**Figure 4D**). This interaction interface is further stabilised by electrostatic interactions between Lys344 and Lys348 of the CL and Asp385 and Asp245 of the C-core, as well as several hydrogen bonds (**Figure 4D, Table S6**). In contrast to the extensive NT and CL-mediated contacts, the area of direct interactions between the rigid C-core domains of adjacent subunits is relatively limited, burying only ∼570 Å^2^ of solvent-accessible surface area. The contacts are mediated by a mixture of polar and hydrophobic interactions (**Figure S8**), including one salt bridge between Asp234 and His320 side chains (**Figure 4C)**.

Comparison with other rhabdoviruses shows substantially lower conservation of residues involved in protein-protein interactions than of residues involved in RNA binding **(Figure 3F**, **Figure 4F**). The NT arm is particularly divergent, with no sequence conservation across the *Rhabdoviridae* family and considerable variability even within the *Ephemerovirus* genus (**Figure S7, Data S1**). For example, Lys7, involved in salt bridge formation with Glu355, is conserved in only five of the thirteen analysed *Ephemerovirus* species. In the remaining species, this residue is substituted by either Gln, Gly, Glu or Asp, eliminating or reversing the charge and preventing salt bridge formation (**Data S1**). Likewise, Lys348, which in BEFV forms a salt bridge with Asp385, is replaced in two ephemeroviruses (Obodhiang and Adelaide River viruses) by serine. Lys348 is also substituted by Cys349 in VSV, and by Ser401 in RABV (**Figure 4D, 4F**).

Interestingly, while the total interaction areas between adjacent C-cores are similar among *Rhabdoviridae* members, molecular interactions between them are generally not conserved. In particular, His320 is substituted by Cys333 in RABV (**Figure 4C, 4F**), and although this region was not resolved in the VSV N structure, sequence alignment indicates His320 substitution by isoleucine. The chemistry of polar/charged residues at C-core interfaces varies among *Rhabdoviridae* members, yet some interactions appear to be conserved. For example, the Asp234/His320 pair in BEFV corresponds to Lys236/Glu2 in VSV, potentially preserving the salt bridge interaction. Several other charged residues at the C-core/C-core interacting region of the BEFV N protein are substituted by hydrophobic residues in other rhabdoviruses. Asp245 corresponds to Thr247 in VSV and Leu258 in RABV, while Glu309 is replaced by Thr311 in VSV and Gln321 in RABV (**Figure 4F, Data S1**). Likewise, Glu319 is substituted by Ala332 in RABV, although a negatively charged residue (aspartic acid) is retained in VSV, as suggested by the sequence alignment (**Figure 4F**).

## Discussion

This study provides the first structural characterisation of a BEFV nucleoprotein-RNA complex, providing a structural basis for understanding genome encapsidation and nucleoprotein assembly of members of the *Ephemerovirus* genus. Despite sharing only relatively low sequence identity with previously characterised *Rhabdoviridae* nucleoproteins, BEFV adopts not only a similar bi-lobed domain organisation, characteristic of the wide *Mononegavirales* order, but also encapsidates RNA along a similar trajectory and with the same 9 nt encapsidation repeat length as seen for VSV [16] and RABV [14,17,19,26,27]. At the same time, the structure unveils distinctive features of the oligomerisation interface that suggest a molecular mechanism by which rhabdoviruses form bullet-shaped nucleocapsids.

### Decameric ring likely nucleates virus assembly

We show that recombinant BEFV N predominantly assembles into decameric rings in the presence of non-specific RNA. Highly similar oligomeric assemblies, also composed of 10 subunits (VSV) or 11 subunits (RABV), have previously been revealed by structural studies of recombinantly produced *Rhabdoviridae* N proteins [14,16,26], but were considered as consequences of crystallisation. Experimental observation of assemblies with essentially identical architecture suggests that ring formation may be an intrinsic property of rhabdovirus nucleoproteins rather than an experimental artefact.

Several observations suggest that the decameric nucleoprotein assembly represents a physiologically relevant nucleation intermediate. First, decameric particles constituted the overwhelming majority of particles observed in this study (91%) (**Figure S3**). Second, the inclination angle of BEFV N subunits with respect to the central axis of 25.96° (**Figure 5A**) closely resembles the ∼25° inclination of subunits within the tip of the bullet-shaped VSV virion [19,25]. Third, the circular architecture of the ring provides a highly curved template, from which successively wider helical turns could emerge. Together, these observations suggest that the decameric ring serves as a nucleation assembly to build the bullet-shaped nucleocapsid.

**Figure 5.**
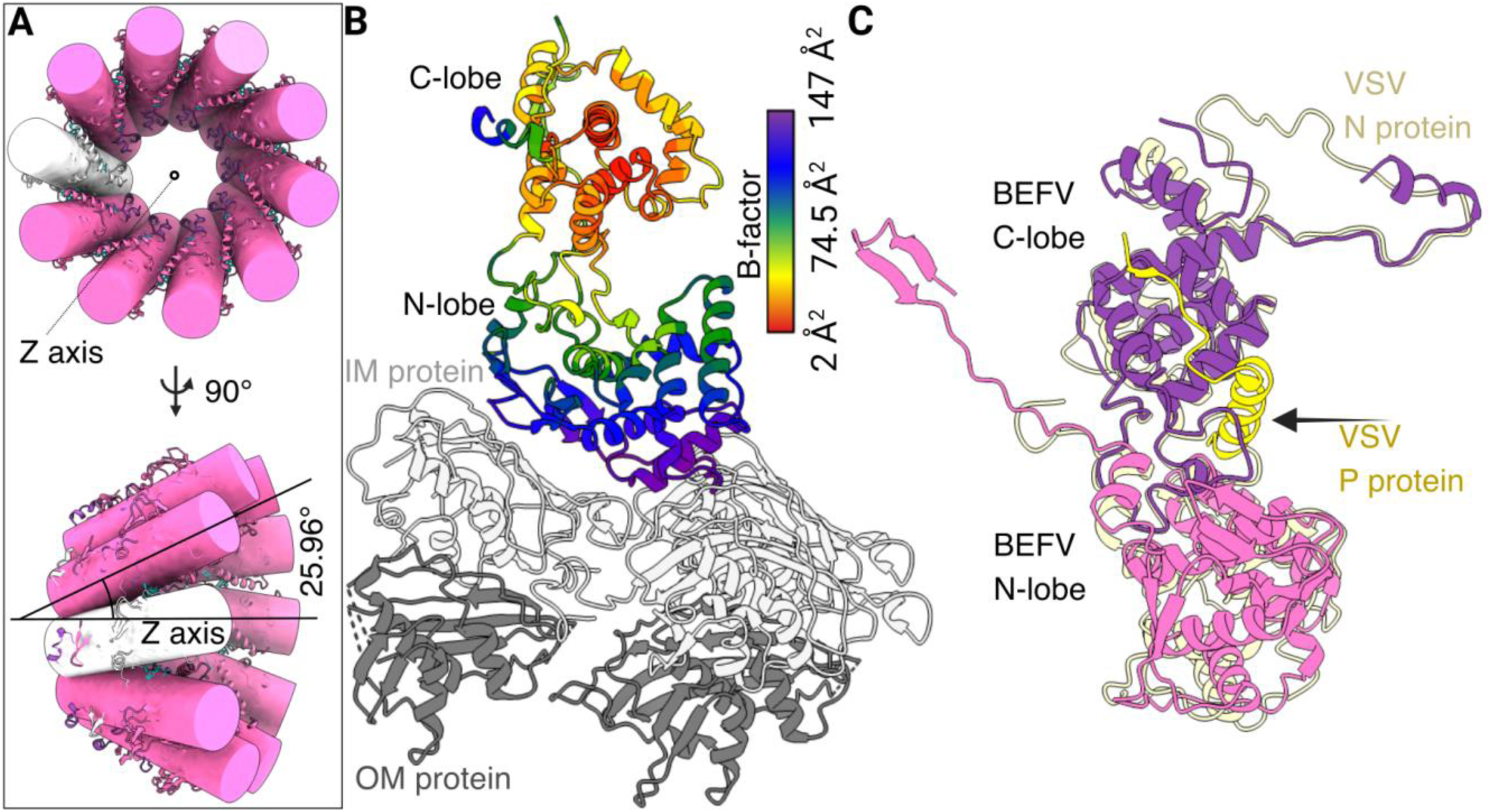
Structural context of BEFV N in virion assembly. (A) Subunit orientations within the decamer. Axial (top) and side (bottom) views with cylinders fitted over ribbon models of individual subunits. The central z-axis and one of the cylinder axes (black lines) were used to calculate the subunit inclination angle. (B) BEFV N-M protein complex, predicted based on the superimposition with the VSV N-M protein assembly (pdb: 7UWS). Inner matrix protein (IM) that interacts with the N-lobe is in light grey, and the outer matrix protein (OM) that interacts with the IM is in dark grey. The N protein of VSV is hidden for clarity. (C) BEFV N-P protein complex, predicted based on superimposition with the structure of the VSV N-P assembly (pdb: 3PMK). The BEFV N ribbon is in purple and pink, the VSV N is in light yellow and the N-terminal part of the VSV P protein in darker yellow.

### Distinct oligomerisation strategy of rhabdoviruses compared to other *Mononegavirales* members

Despite extensive sequence variation, the overall organisation of the N-RNA complex assembly is highly conserved among the *Rhabdoviridae* family. Of particular significance is the combination of the relatively small C-core/C-core interface with extensive interactions mediated by the NT arm and CL, which are flexibly connected to the main bodies of subunits. As the C-core/C-core interface is the only direct contact area between the main bodies of subunits, it is likely to define their relative orientation. Its limited size and relatively flat geometry are consistent with a pivot-like interface that can facilitate rigid-body rotation between adjacent subunits. These positional adjustments can be accommodated by repositioning the flexible NT/CL regions, maintaining the integrity of the assembly. Such subunit-subunit interactions could serve as a structural basis for progressive changes in the assembly’s curvature, required to generate a bullet-shaped nucleocapsid. In contrast, nucleoprotein assemblies of other members of *Mononegavirale*s, which form nucleocapsids of a constant radius, such as Ebola, are stabilised through extensive interaction surfaces between the globular domains of neighbouring subunits [20] rather than by flexible terminal extensions. This architecture generates a comparatively rigid assembly with a constant filament diameter. The apparent flexibility in subunit-subunit interactions of rhabdoviruses may therefore represent a specific adaptation enabling curvature variation during nucleocapsid assembly.

In the BEFV N protein, five regions correspond to diffused cryo-EM density with elevated B-factor values (**Figure 1D, Figure S4**), indicating their flexibility. Interestingly, intrinsically disordered regions were previously predicted in the BEFV nucleoprotein based on a sequence similarity analysis [49], matching all these regions. Four of these regions are located on the distal part of the subunit in the N-lobe region. In the mature virion, these flexible regions are likely important for mediating interactions with the matrix protein (**Figure 5B**), in common with analogously positioned flexible loops identified in the VSV N-M protein complex [19,25] (**Figure 5B**).

The C-lobe of BEFV N protein contains one unmodelled disordered region corresponding to residues 355-377, forming a part of the flexible loop. The position of this loop structurally corresponds to the observed C-loop, which includes residues 341-374 in VSV nucleoprotein [17,19]. It plays a role in mediating contact with the phosphoprotein (P protein), which is associated with the L protein (polymerase), a vital component for genome replication [12,26,27,50–52]. The loop is located towards the inner part of the whole virion, becoming ordered after the interaction with the C-terminal part of one out of two P protein subunits in the functional dimer [12,25,49,50]. Based on the functional conservation of P proteins, we suggest the C-loop in the BEFV plays the same role (**Figure S9**).

### *Rhabdoviridae* N proteins share a ‘3-out, 1-in, 1-out, 2-in, 2-out’ conformation of bound RNA

The studied conformation of the BEFV RNA was shown to be similar to that of the VSV [16,19,25] and RABV [14,26], suggesting that this feature is conserved within the whole *Rhabdoviridae* family. However, this conformation is not shared within the wider *Mononegavirales* order, where all *Filoviridae, Paramyxoviridae* and *Pneumoviridae* members showcase ‘3-in, 3-out’ conformation (**Figure 3E**) [20–23].

As binding of the -ssRNA genome to the nucleoprotein is mainly mediated by basic amino acids and is non-specific across the order, representatives of the *Mononegavirales* order require strict control of genome recognition and virion assembly by other factors [12,50]. In the archetypal species of the *Rhabdoviridae* family, VSV and RABV, the P protein has been shown to guide nucleoprotein-RNA assembly during viral replication [17,18]. In VSV, the N-terminal part of the P protein, consisting of residues 1-60, extends into the RNA-binding cleft, filling it and thus preventing binding to a random cellular RNA (**Figure 5C**). Moreover, it competes with the N-terminal arm of the N protein, thereby modifying how N protein subunits interact with each other (**Figure S9**) [12,50]. To analyse the possibility of a similar strategy occurring in the BEFV virion, the VSV N-P structure was superimposed with BEFV N (**Figure 5C**). The overlay showed that the N-terminus of P accommodates the same position within the BEFV and VSV N subunits [12,50]. The same strategy is likely used in the rest of the *Ephemerovirus* genus, as they show a high sequence conservation in the RNA-binding groove (**Figure S7**).

### Mechanism of the bullet-shaped virion assembly

Based on our findings, we propose the following mechanism for bullet-shaped virion assembly (**Figure 6**). Upon viral infection, the L enzyme, comprising RNA-dependent RNA polymerase (RdRP) activity, is activated in the host cytosol, initiating transcription of the viral genome. Newly synthesised N protein subunits interact with the N-terminal arm of the P protein, which temporarily blocks RNA binding [12]. The genome is recruited to N subunits by a single P protein subunit associated with both L and N proteins [12,53]. As the RNA strand is produced, new N subunits are added to the complex, replacing the P protein [12,53]. We propose that this process initially generates a decameric N-RNA assembly that forms the tip of the future virion. Subsequent incorporation of additional N subunits, guided by the growing genomic RNA chain, is enabled by subtle changes in the intersubunit orientation, facilitated by the flexible NT and CL regions. The small C-core/C-core interface serves as a pivot for these adjustments, allowing successive expansion of the assembly through the formation of wider helical turns. As the elongation proceeds, the N-RNA complex grows into a wider helical structure, with the number of N subunits per turn increasing progressively, for example, to ∼33–42 in VSV [19,25].

**Figure 6.**
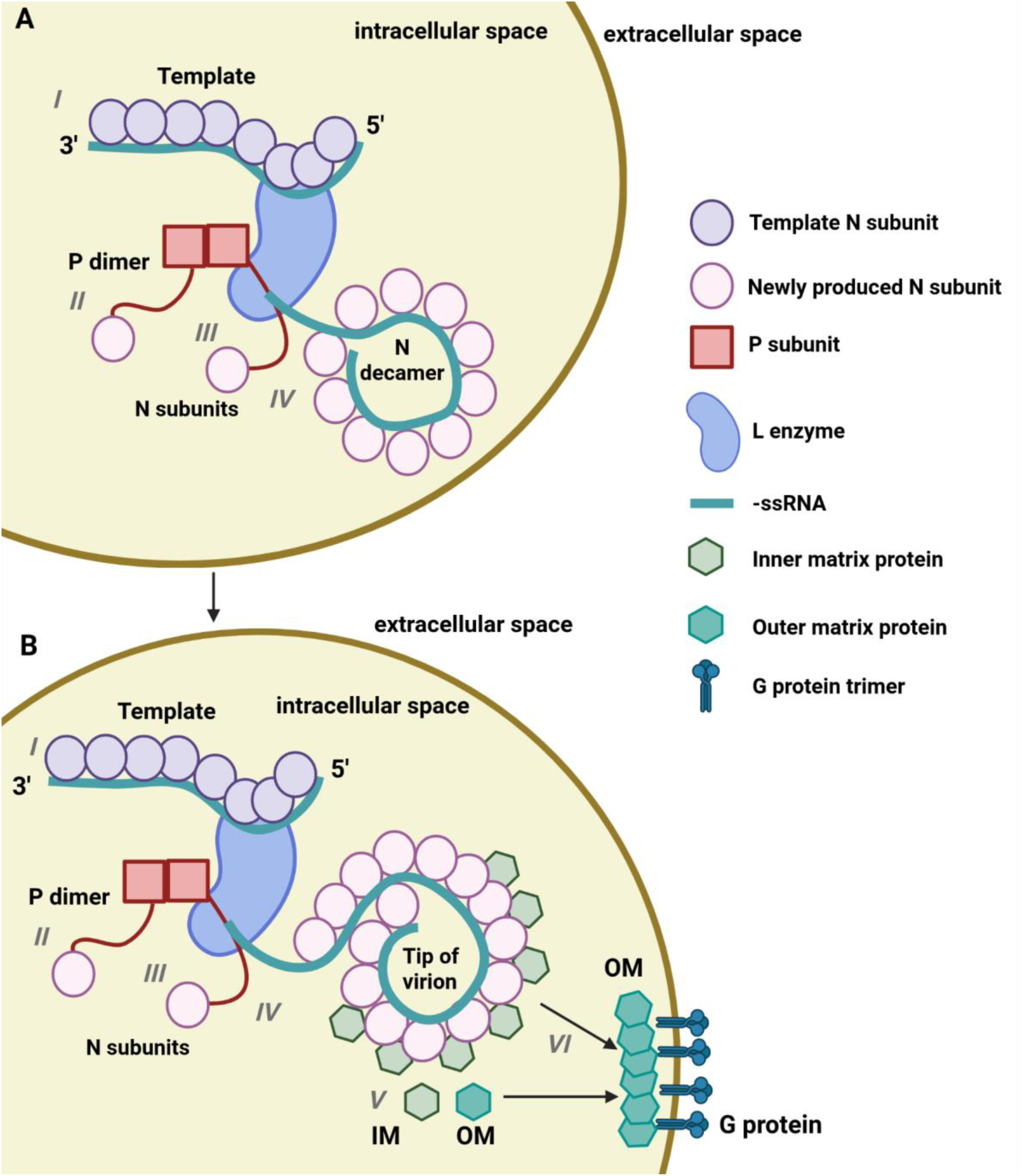
Proposed model of the BEFV virion assembly. (I) The ribonucleoprotein complex of the entered BEFV virion is used as a template for transcription and replication by L, comprising RNA polymerase activity. (II) P protein dimer binds to both L and newly translated N proteins, preventing non-specific RNA binding. (III) One P subunit simultaneously interacts with L, a new N subunit and newly synthesised negative-sense genomic RNA. (IV) P mediates a transfer of new N subunits onto the growing RNA chains synthesised by L. Following release from P, N proteins form the decameric assembly with RNA, which serves as the tip, nucleating formation of the future virion. (V) Newly produced inner matrix protein subunits (IM) bind to the N-RNA complex, promoting the shift of the decameric assembly to helical, and stabilising the helical nucleocapsid assembly. (VI) The assembled N-RNA-IM complex is transported to the plasma membrane, where outer matrix protein (OM) and glycoprotein (G) assemblies will facilitate virion maturation and eventual budding from the host cell (created in https://BioRender.com).

Interaction with the inner matrix protein (IM) likely stabilises helical assembly and facilitates the transition of the initial ring into the helical structure [19,25]. The packaged N-RNA complex, together with IM, is delivered to the host cell’s plasma membrane, where other structural proteins, such as outer matrix protein (OM) and glycoproteins (G), accumulate [10,53]. G proteins interact with the OM, which connects to the IM-N-RNA assembly. As a result, the virion matures, and the new progeny of the virus exit the cell via budding.

In summary, the BEFV nucleoprotein structure and its comparison with other negative-sense RNA viruses identify structural principles that may define the distinctive bullet-shaped morphology of rhabdoviruses. A small C-core/C-core interface combined with NT- and CL-mediated tethering interactions appears well suited to accommodate progressive changes in curvature during nucleocapsid assembly, while maintaining structural integrity and continuous genome encapsidation. Whereas RNA recognition is strongly conserved across the *Rhabdoviridae* family, oligomerisation interfaces exhibit substantial plasticity, indicating that evolutionary constraints act primarily to preserve assembly architecture rather than individual residue interactions. These findings provide a structural basis for understanding rhabdovirus assembly and may inform the development of novel antiviral strategies.

## Acknowledgements

We thank Sam Hart and Johan Turkenburg for their assistance with cryo-EM data collection. We also acknowledge the Technology Facility at the Biology Department of the University of York for access to instrumentation and technical support. We thank Maria Chechik for laboratory support and Huw Jenkins for advice in computational analysis. We also thank Chris Hill for insightful discussions and advice.

