## Supplementary Information for "Cryo-electron microscopy structure of the bovine ephemeral fever virus RNA-nucleoprotein assembly"

**This file includes:**
Figures S1 to S9
Tables S1 to S6
Supplementary References 1 to 3

**Other supporting materials include the following:**

**Data S1.** Phylogenetic tree and multiple sequence alignment based on nucleoprotein sequences of *Mononegavirales* species.

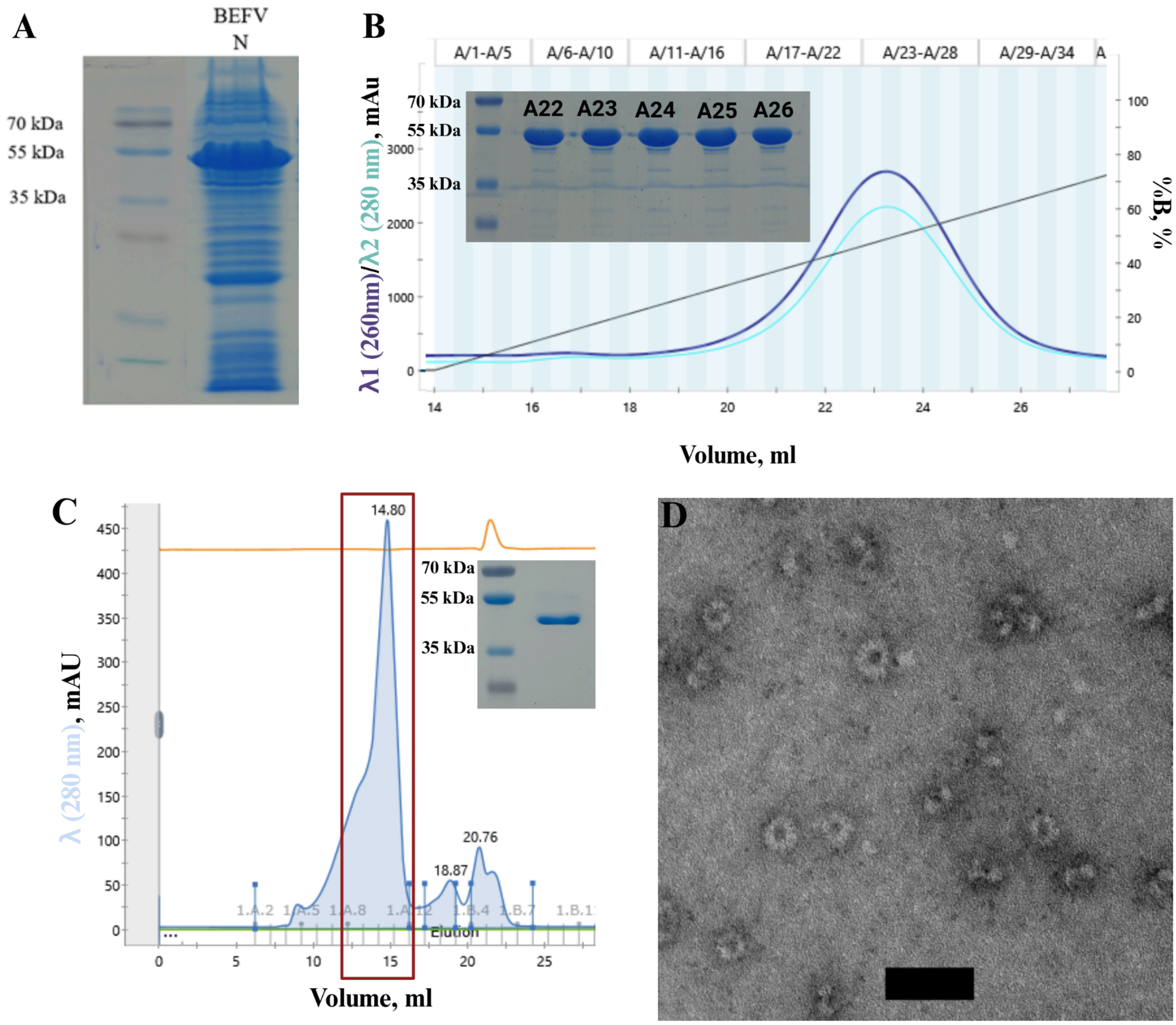


**Figure S1**. **Production and purification of BEFV nucleoprotein.**
(A) SDS-PAGE electrophoresis of the nucleoprotein after production in *E.coli* showing abundant protein of ~52 kDa size, which corresponds to the expected size. (B) Chromatogram of the nucleoprotein obtained using immobilised metal affinity chromatography with SDS-PAGE of phases A22-A26 shown in the left corner. The absorbance in milli-absorbance units (mAU) at 260 nm wavelength is represented in blue and at 280 nm in cyan. The right y-axis shows the concentration of the elution buffer, depicted in a black line. (C) Chromatogram of BEFV nucleoprotein obtained using the size exclusion chromatography: x axis – absorbance (mAU); y axis – volume (ml), 1.A.2 – 1.C.6 – rack/tube positions; blue curve – absorbance at 280 nm. Numbers above peaks show the elution volume of their maxima. The collected fractions are highlighted in a red rectangle. The SDS-PAGE of the pooled sample is shown at the top right corner. (D) Negative stain TEM image of BEFV nucleoproteins at 150,000x magnification, scale bar 50 nm.

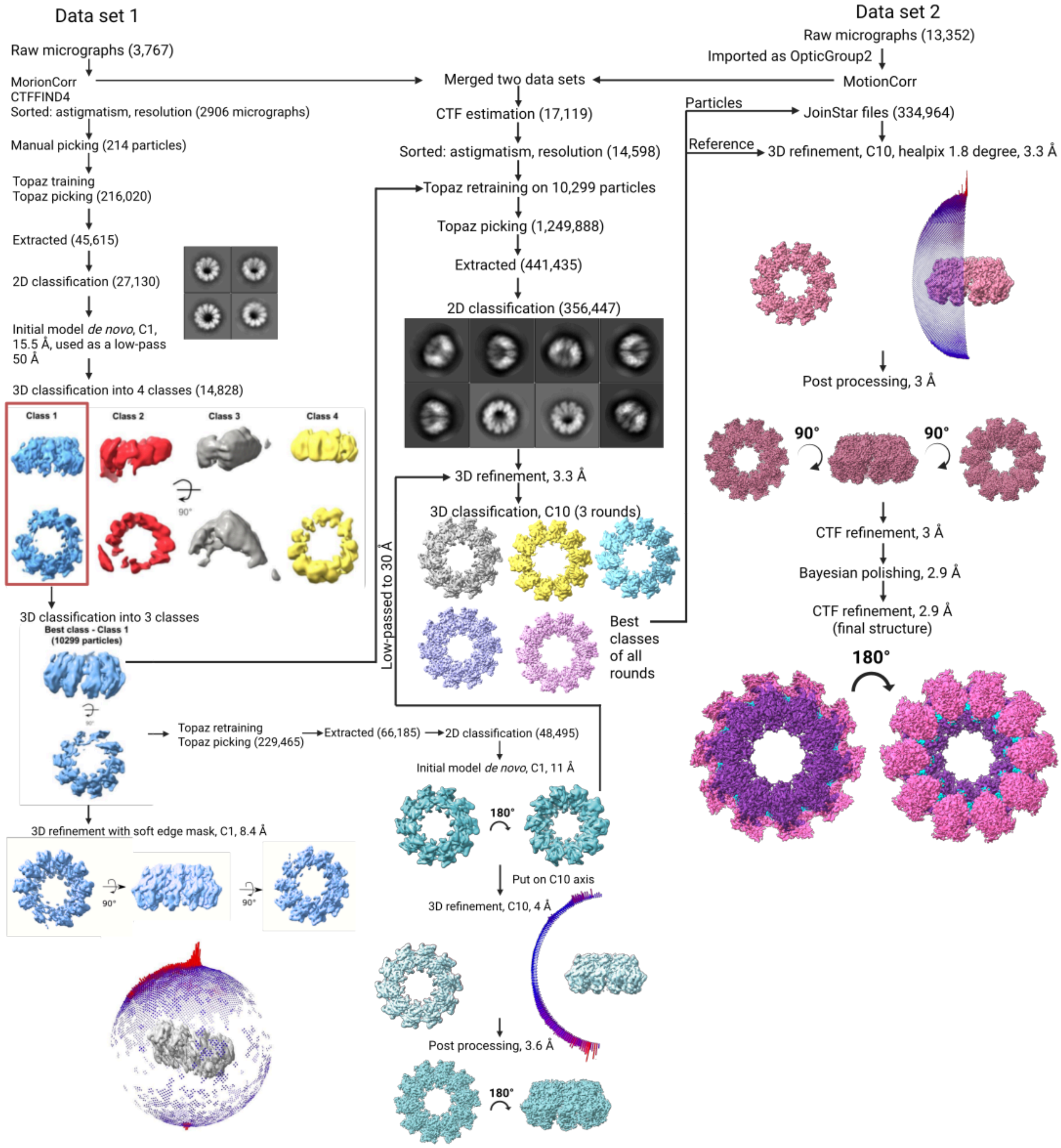


**Figure S2**. **Cryo-EM single-particle analysis (SPA) workflow of the nucleoprotein.** The blue-and-red spheres around maps represent Euler angular distribution plots, which represent the particles' orientation distribution. The colouring from blue to red represents the number of particles of the specific orientation, with red showing the largest number of particles in that orientation. The gaps mean the absence of a specific orientation in the dataset.

A

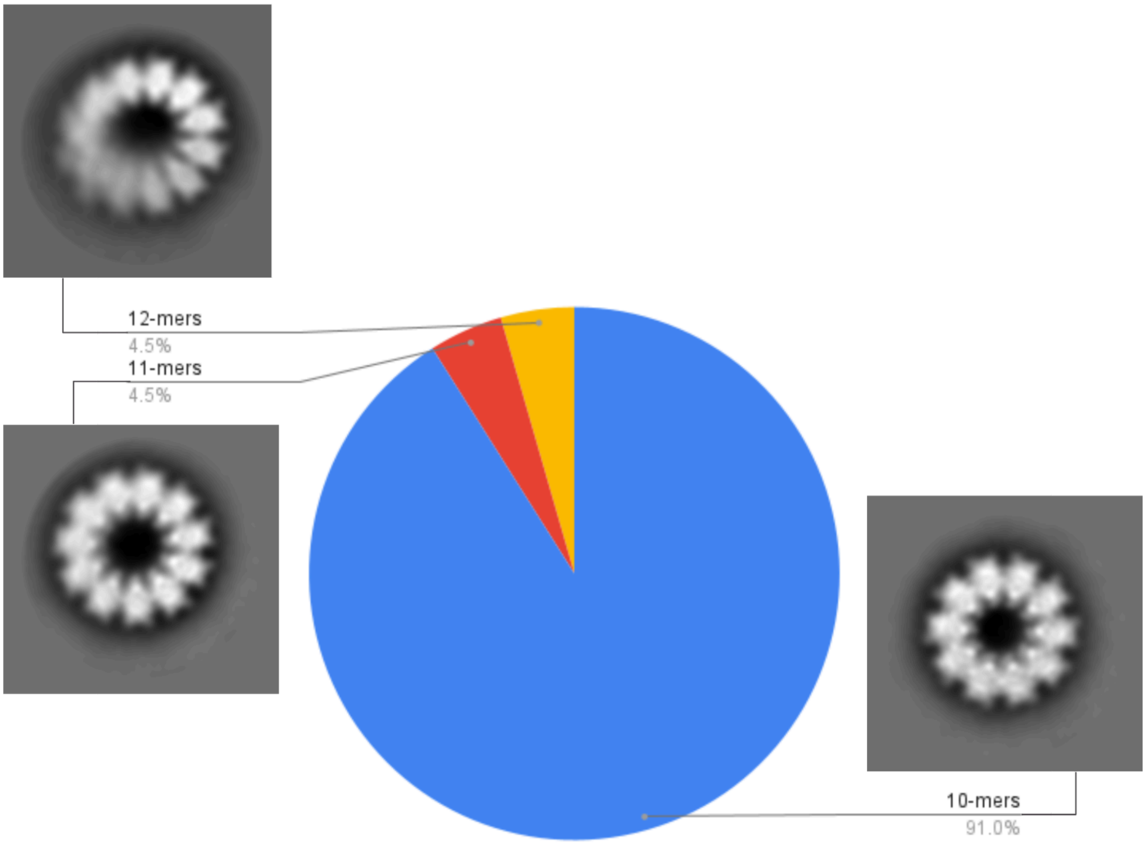


B

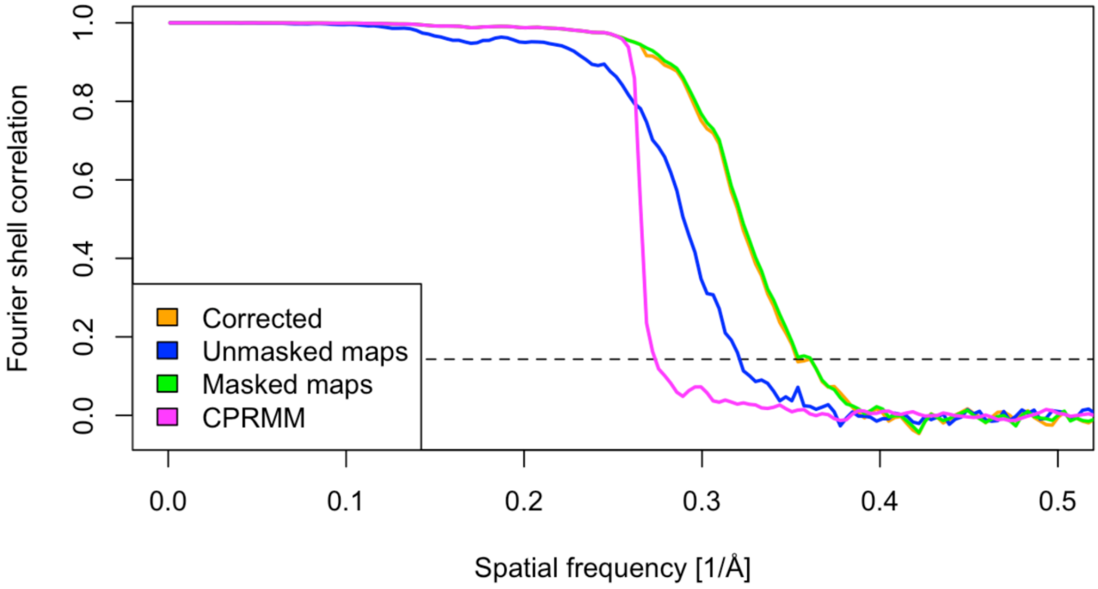


**Figure S3. BEFV N data analysis.**
(A) A pie chart showing the number of subunits per ring in a processed BEFV N protein dataset, as calculated from addressable 2D class averages. (B) FSC graph for the final BEFV N structure with the threshold of 0.143 used for the resolution estimation shown in the dashed black line.

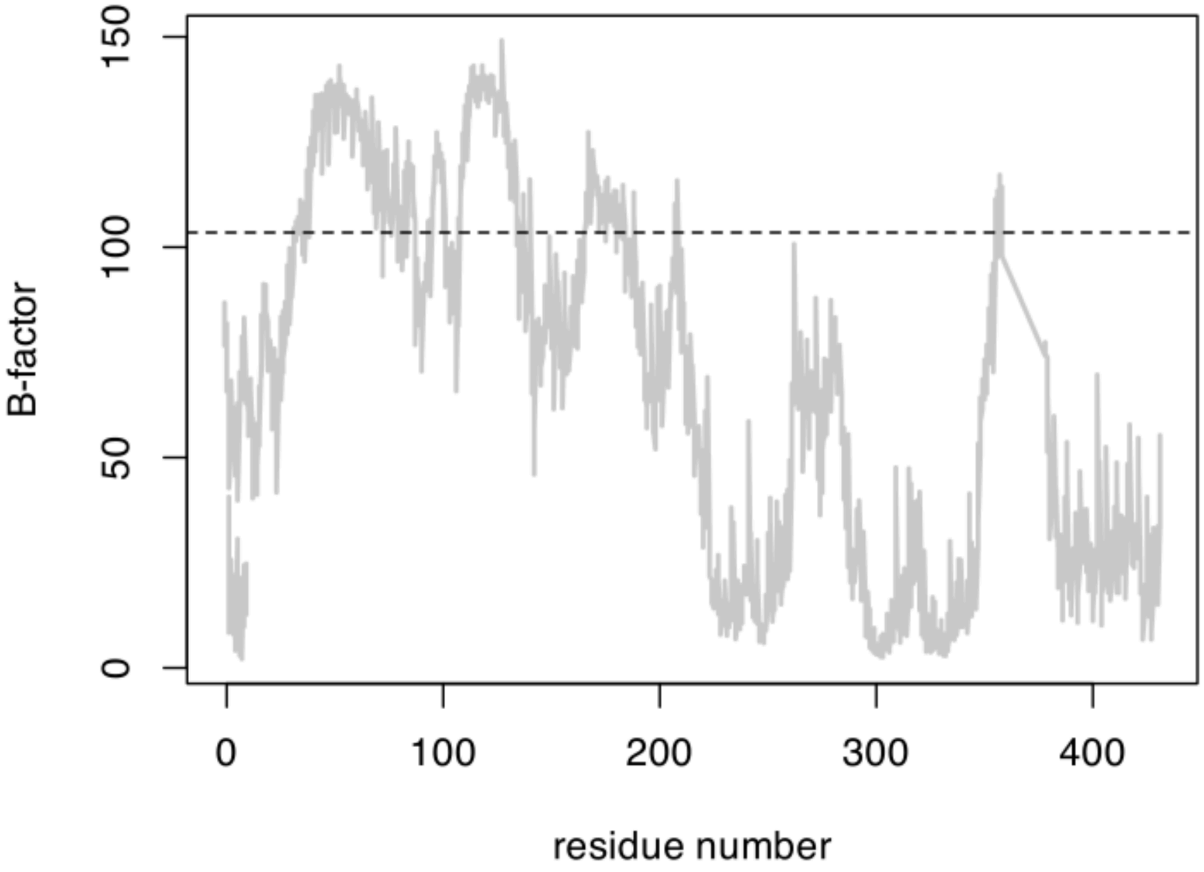


**Figure S4. B-factor plot of BEFV N model.**
A threshold is shown with a dashed line, representing the value at which density becomes diffused. Residues with high B-factor values (above the dashed threshold) cannot be used to confidently refine the *de novo* model, and the AlphaFold3-predicted [1] model was required to guide the model building.

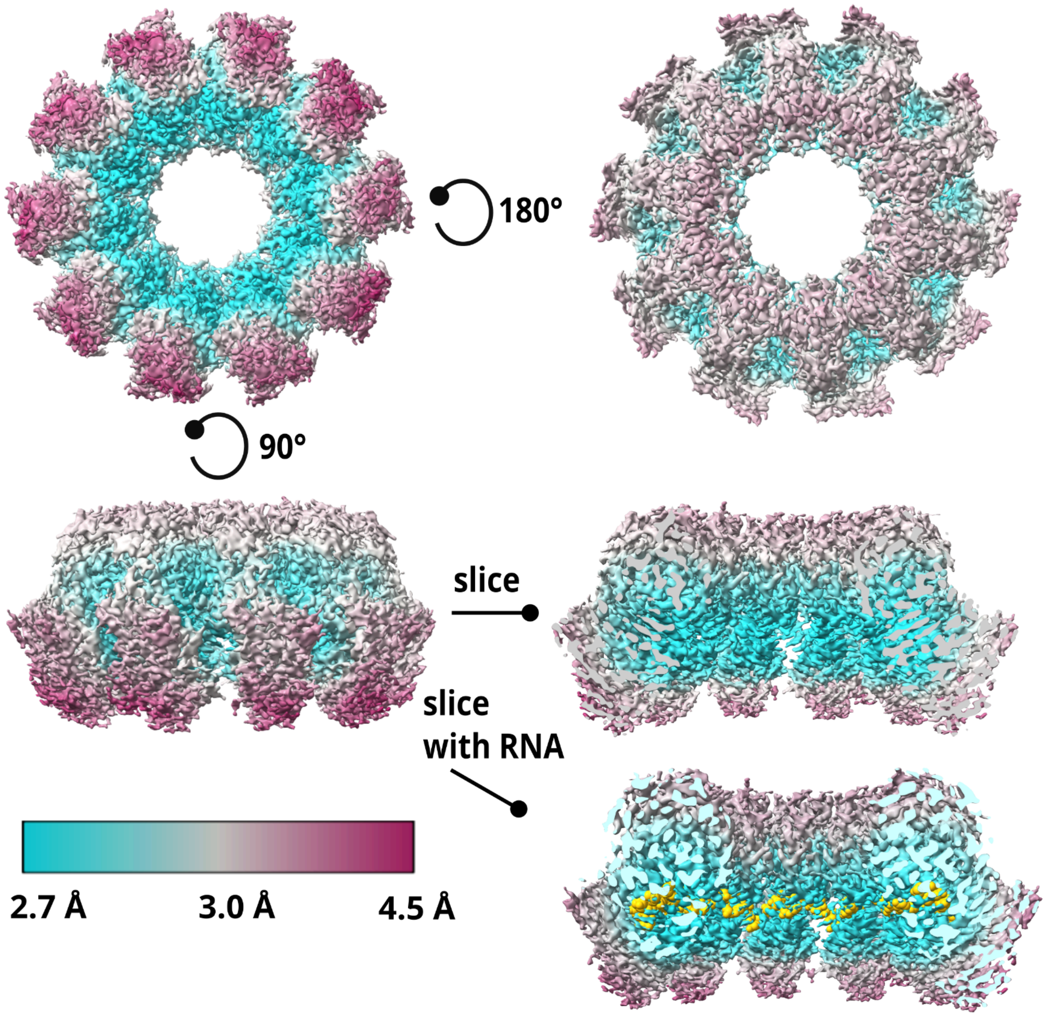


**Figure S5. Local resolution.**
Local resolution of the nucleoprotein cryo-EM density map (left) and the same map sliced from the centre (right), represented in colour coding from 2.7 Å to 4.5 Å. At the bottom, the ssRNA model is showcased in the sliced map as yellow spheres.

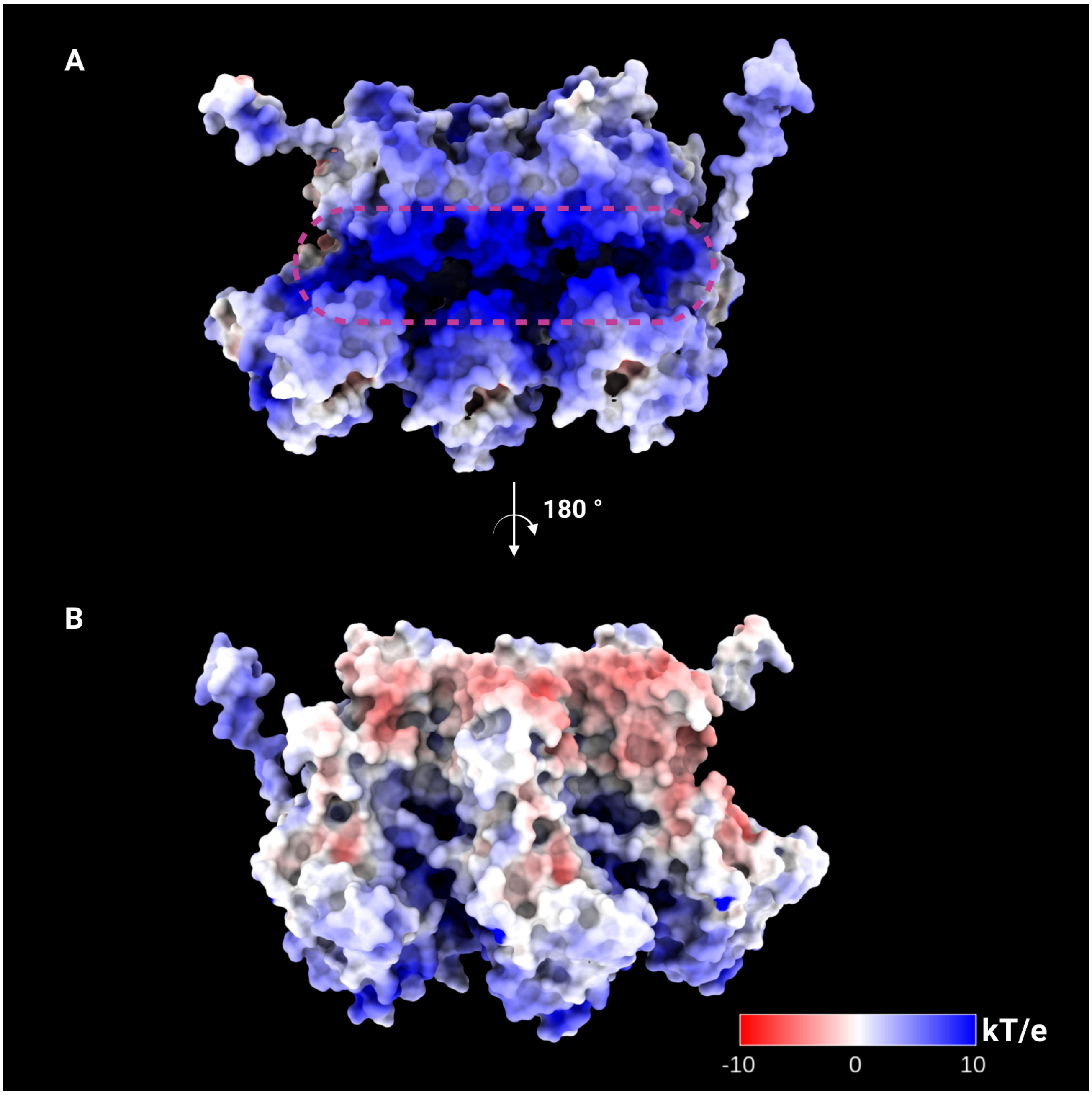


**Figure S6. Electrostatic potential map of BEFV N three subunits.**
Molecular surface corresponding to three BEFV protein subunits coloured according to the electrostatic potential. The most positively charged area corresponds to the N protein-ssRNA binding interface. (A) The surface area is viewed from the inside of the ring. The RNA-binding region is outlined with a dashed magenta oval. (B) The surface area is viewed from the outside of the ring.

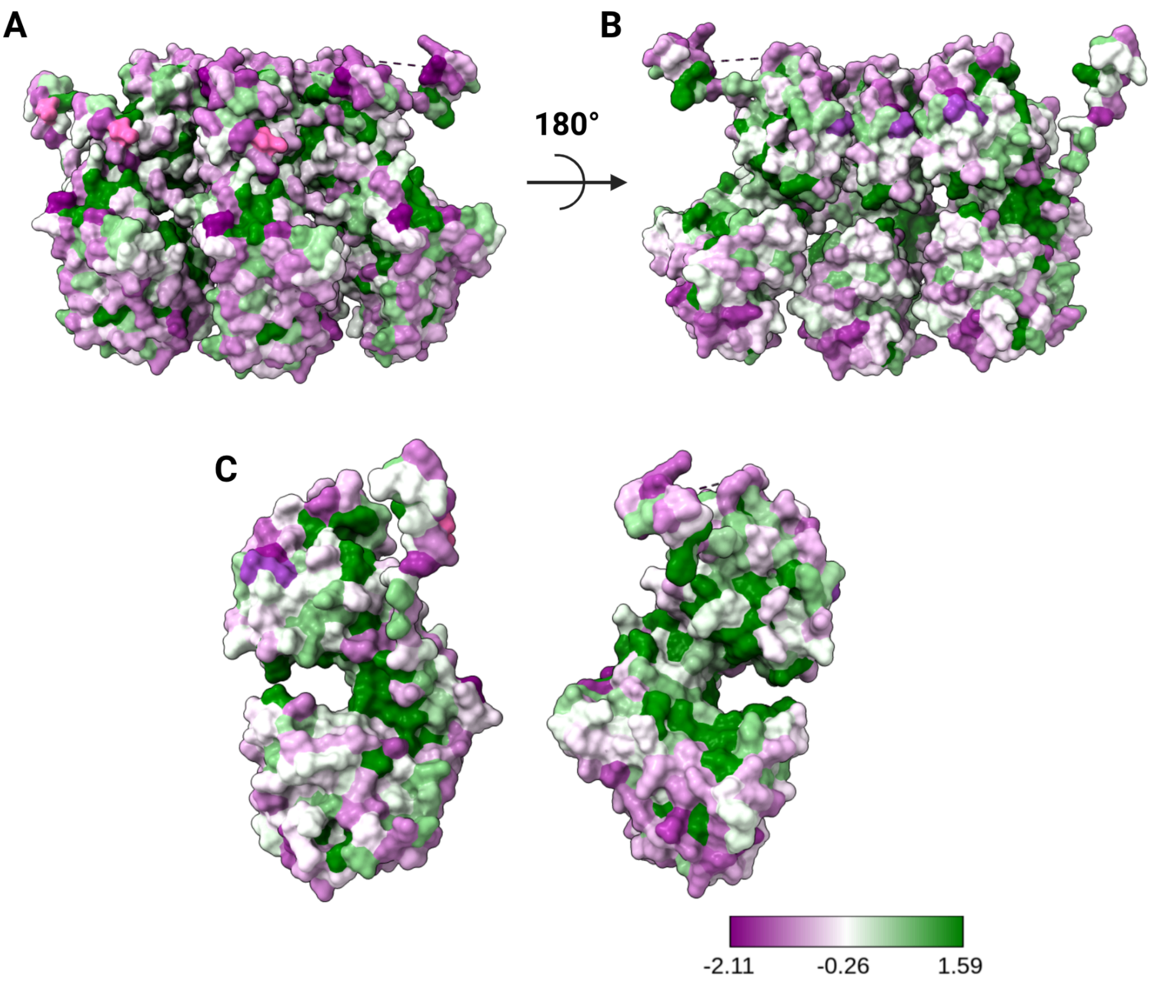


**Figure S7. Sequence conservation of the *Ephemerovirus* nucleoproteins mapped on BEFV N.**

(A) Surface representation of three subunits of the BEFV N coloured based on sequence conservation, where the green surface represents the most conserved areas, and purple represents the least conserved areas. (B) Surface representation of three subunits of the BEFV N rotated 180° with the RNA-binding cleft pointing towards the front. (C) Two neighbouring subunits rotated 180° from each other, showcasing the sequence identity of their interaction zones.

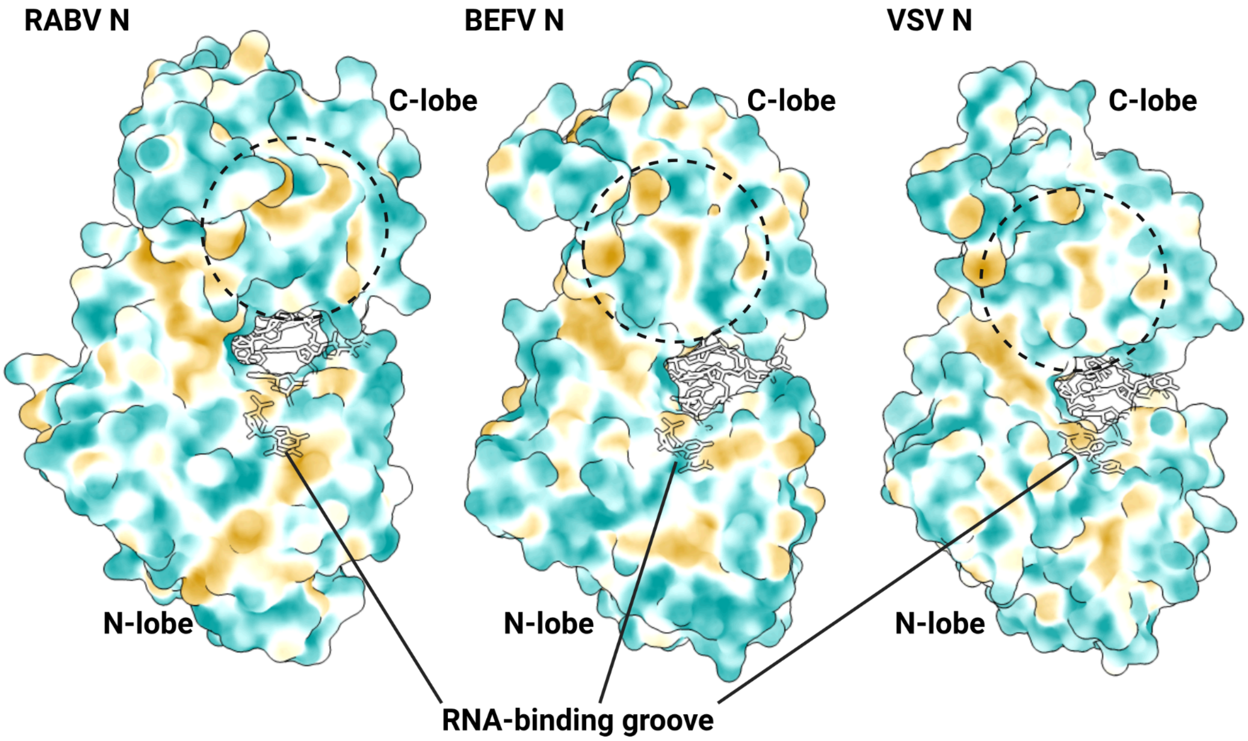


**Figure S8. Hydrophobicity representation of RABV N protein subunit, BEFV N and VSV N with the emphasis on RNA-binding grooves and C-lobes.** The subunits are shown with the RNA-binding groove pointing to the right. RNAs are shown as transparent sticks. The C-lobe/C-lobe interaction area is highlighted in a black dashed circle. Blue patches on N protein represent a polar region; yellow patches represent a hydrophobic region; and white patches represent an amphipathic region.

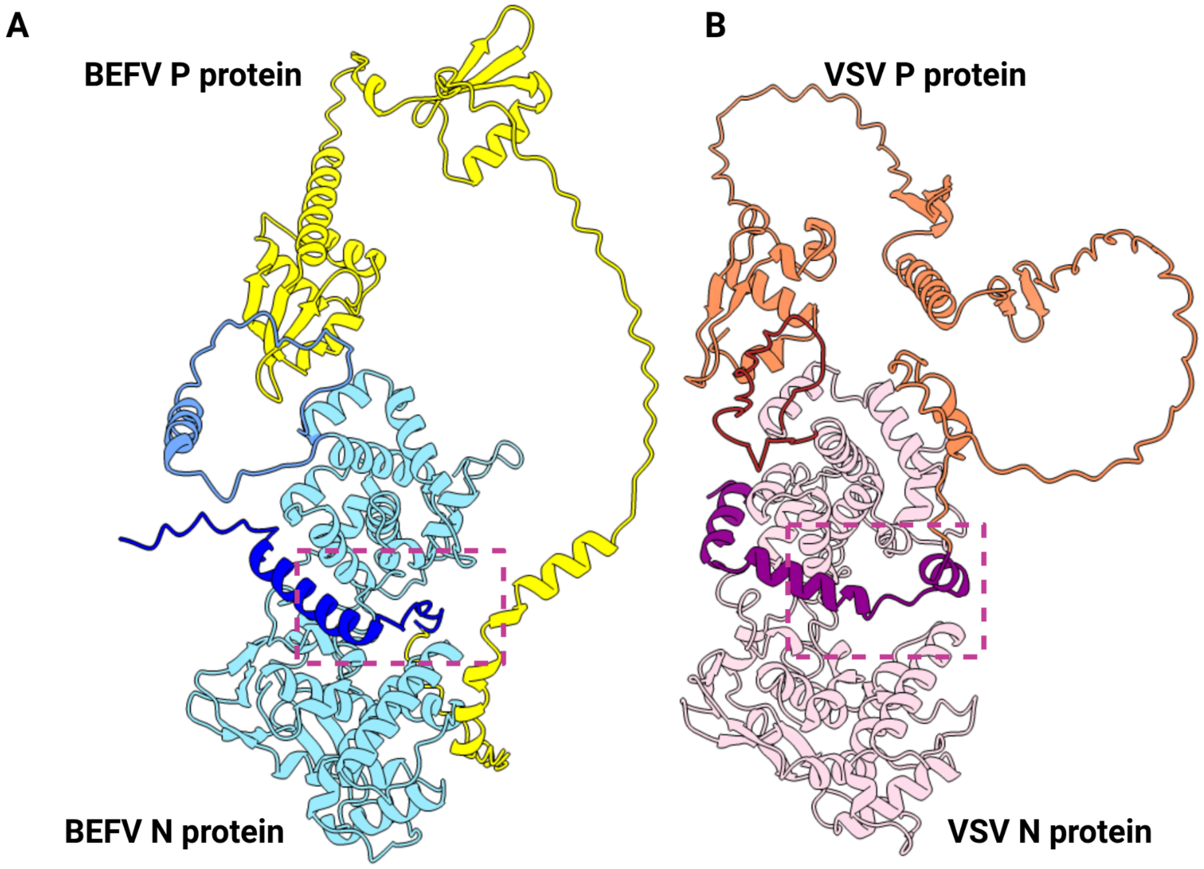


**Figure S9. AlphaFold3 predicted models for VSV N-P and BEFV N-P.** (A) Predicted model of BEFV N-P complex with one N subunit shown in blue and one P subunit shown in yellow. The N-terminus of the BEFV P protein binding to the RNA groove is shown in dark blue. The BEFV nucleoprotein C-loop that is stabilised by the P protein is depicted in a different shade of blue. (B) Predicted model of VSV N-P complex with one N subunit shown in light pink and one P subunit shown in coral. The N-terminus of the VSV P binding to the RNA groove is shown in purple. The VSV nucleoprotein C-loop that is stabilised by the P protein is depicted in brown. (A-B) Magenta dashed boxes showcase the RNA-binding groove.

**Table S1. Data collection and image processing statistics.**

| | Data set 1 | Data set 2 |
|---|---|---|
| **Data collection** | | |
| Grids type | UltraAuFoil R1.2/1.3 | UltraAuFoil R1.2/1.3 with 0.01% poly-l-lysine |
| Magnification | ×240,000 | ×240,000 |
| Pixel size (Å) | 0.574 | 0.574 |
| Electron exposure (e-/$Å^2$) | 50 | 50 |
| Exposure time (sec) | 3.41 | 2.68 |
| Defocus range (µm) | -0.6 to -2.0 with step 0.2 | -0.6 to -1.6 with step 0.2 |
| Voltage (kV) | 200 | 200 |
| Detector | Falcon 4 | Falcon 4 |
| Microscope | Glacios (TFS), York | Glacios (TFS), York |
| **Data processing** | | |
| Processing software | RELION5 | |
| Symmetry | C10 | |
| Initial number of particles | 441,435 | |
| Final number of particles | 334,964 | |
| Initial model | *de novo* | |
| Map resolution (Å) | 2.9 | |
| Sharpening B factor ($Å^2$) | -93.4 | |

**Table S2. Model building, refinement and validation statistics.**

| Parameters | Statistics |
|---|---|
| **Model composition** | |
| Non-hydrogen atoms | 3322 |
| Protein residues | 414 |
| Nucleic acid bases | 9 |
| **Validation** | |
| Ramachandran Favoured (%) | 99% |
| Ramachandran Outliers | 0 |
| Rama-Z (Z-score, RMSD) | -1.12 (0.37) |
| Rotamer outliers | 0.8% |
| Clashscore | 3 |
| Molprobity score | 1.11 |
| CaBLAM outliers | 0.74% |
| Q-score | 0.564 |
| PDB entry | 9U0F |

**Table S3**. **BEFV genomic RNA sequence alignment analysis.**
Sequence grouped by every 9 nucleotides and represented as letter Y for pyrimidines and R for purines according to IUPAC nomenclature, showing their ~60/40 distribution over the sequence.

| Start nucleotide number | BEFV genome sequence representation, from 3` to 5` | Pyrimidines (Y)/purines (R) BEFV genome sequence representation, from 3` to 5` | Percentage of pyrimidines (Y) in a position | Percentage of purines (R) in a position |
|---|---|---|---|---|
| 1 | [UGCUCUUUU…] | [YRYYYYYYY…] | 1 - 55.62%<br>2 - 59.78%<br>3 - 58.51%<br>4 - 57.49%<br>5 - 61.41%<br>6 - 57.43%<br>7 - 54.95%<br>8 - 60.45%<br>9 - 57.31% | 1 - 44.38%<br>2 - 40.22%<br>3 - 41.49%<br>4 - 42.51%<br>5 - 38.59%<br>6 - 42.51%<br>7 - 44.99%<br>8 - 49.49%<br>9 - 42.63% |
| 2 | U[GCUCUUUU…] | Y[RYYYYYYY…] | 1 - 59.78%<br>2 - 58.51%<br>3 - 57.49%<br>4 - 61.41%<br>5 - 57.43%<br>6 - 54.95%<br>7 - 60.45%<br>8 - 57.31%<br>9 - 55.56% | 1 - 40.22%<br>2 - 41.49%<br>3 - 42.51%<br>4 - 38.59%<br>5 - 42.51%<br>6 - 44.99%<br>7 - 39.49%<br>8 - 42.63%<br>9 - 44.38% |
| 3 | UG[CUCUUUU…] | YR[YYYYYYY…] | 1 - 58.51%<br>2 - 57.49%<br>3 - 61.41%<br>4 - 57.43%<br>5 - 54.95%<br>6 - 60.45%<br>7 - 57.31%<br>8 - 55.56%<br>9 - 59.78% | 1 - 41.49%<br>2 - 42.51%<br>3 - 38.59%<br>4 - 42.51%<br>5 - 44.99%<br>6 - 39.49%<br>7 - 42.63%<br>8 - 44.38%<br>9 - 40.16% |
| 4 | UGC[UCUUUU…] | YRY[YYYYYY…] | 1 - 57.49%<br>2 - 61.41%<br>3 - 57.43%<br>4 - 54.95%<br>5 - 60.45%<br>6 - 57.31%<br>7 - 55.56%<br>8 - 59.78%<br>9 - 58.45% | 1 - 42.51%<br>2 - 38.59%<br>3 - 42.51%<br>4 - 44.99%<br>5 - 39.49%<br>6 - 42.63%<br>7 - 44.38%<br>8 - 40.16%<br>9 - 41.49% |
| 5 | UGCU[CUUUU…] | YRYY[YYYYY…] | 1 - 61.41%<br>2 - 57.43%<br>3 - 54.95% | 1 - 38.59%<br>2 - 42.51%<br>3 - 44.99% |

| | | | | |
|---|---|---|---|---|
| | | | 4 - 60.45%<br>5 - 57.31%<br>6 - 55.56%<br>7 - 59.78%<br>8 - 58.45%<br>9 - 57.43% | 4 - 39.49%<br>5 - 42.63%<br>6 - 44.38%<br>7 - 40.16%<br>8 - 41.49%<br>9 - 42.51% |
| 6 | UGCUC[UUUU…] | YRYYY[YYYY…] | 1 - 57.46%<br>2 - 54.98%<br>3 - 60.48%<br>4 - 57.34%<br>5 - 55.59%<br>6 - 59.82%<br>7 - 58.49%<br>8 - 57.46%<br>9 - 61.39% | 1 - 42.54%<br>2 - 45.02%<br>3 - 39.52%<br>4 - 42.66%<br>5 - 44.41%<br>6 - 40.18%<br>7 - 41.51%<br>8 - 42.54%<br>9 - 38.61% |
| 7 | UGCUCU[UUU…] | YRYYYY[YYY…] | 1 - 54.98%<br>2 - 60.48%<br>3 - 57.34%<br>4 - 55.59%<br>5 - 59.82%<br>6 - 58.49%<br>7 - 57.46%<br>8 - 61.39%<br>9 - 57.40% | 1 - 45.02%<br>2 - 39.52%<br>3 - 42.66%<br>4 - 44.41%<br>5 - 40.18%<br>6 - 41.51%<br>7 - 42.54%<br>8 - 38.61%<br>9 - 42.54% |
| 8 | UGCUCUU[UU…] | YRYYYYY[YY…] | 1 - 60.48%<br>2 - 57.34%<br>3 - 55.59%<br>4 - 59.82%<br>5 - 58.49%<br>6 - 57.46%<br>7 - 61.39%<br>8 - 57.4%<br>9 - 54.92% | 1 - 39.52%<br>2 - 42.66%<br>3 - 44.41%<br>4 - 40.18%<br>5 - 41.51%<br>6 - 42.52%<br>7 - 38.61%<br>8 - 42.54%<br>9 - 45.02% |
| 9 | UGCUCUUU[U…] | YRYYYYYY[Y…] | 1 - 57.34%<br>2 - 55.59%<br>3 - 59.82%<br>4 - 58.49%<br>5 - 57.46%<br>6 - 61.39%<br>7 - 57.4%<br>8 - 54.92%<br>9 - 60.42% | 1 - 42.66%<br>2 - 44.41%<br>3 - 40.18%<br>4 - 41.51%<br>5 - 42.54%<br>6 - 38.61%<br>7 - 42.54%<br>8 - 45.02%<br>9 - 39.52% |

*[] - indicates included in analysis nucleotides

**Table S4. Details on the atoms of residues involved in protein-RNA interactions, forming H-bonds and ionic interactions.**
Estimated using PDBePisa [2,3].

| Number of the contact | RNA (poly-U) | | Distance (Å) | N subunit | |
|---|---|---|---|---|---|
| | Base | Atom | | Residue | Atom |
| 1 | U1 | OP1 | 3.58 | Lys284 | NZ |
| 2 | U2 | OP1 | 2.94 | Ser285 | N |
| 3 | U2 | OP2 | 3.17 | Lys284 | NZ |
| 4 | U3 | OP1 | 2.92 | Ala224 | N |
| 5 | U3 | OP1 | 3.13 | Ala223 | N |
| 6 | U3 | OP2 | 2.89 | Ser289 | N |
| 7 | U3 | OP2 | 2.73 | Ser289 | OG |
| 8 | U3 | O2 | 2.91 | Tyr296 | OH |
| 9 | U4 | OP2 | 3.12 | Arg310 | NH1 |
| 10 | U4 | OP2 | 2.95 | Arg310 | NH2 |
| 12 | U4 | OP2 | 2.77 | Tyr296 | OH |
| 13 | U4 | O4 | 3.12 | Arg310 | NH1 |
| 14 | U4 | O3 | 3.23 | Arg419 | NH1 |
| 15 | U4 | O2 | 3.68 | His313 | ND1 |
| 16 | U4 | O2 | 2.97 | Arg419 | NH1 |
| 17 | U5 | OP1 | 3.59 | Arg419 | NH1 |
| 18 | U5 | O2 | 3.62 | Asn150 | ND2 |
| 19 | U6 | OP1 | 3.45 | Arg419 | NH1 |
| 20 | U6 | OP1 | 3.08 | Arg419 | NH2 |
| 21 | U7 | OP1 | 3.75 | Lys158 | NZ |
| 22 | U7 | OP2 | 3.18 | Arg142 | NH2 |
| 23 | U7 | OP2 | 3.46 | His151 | NE2 |
| 24 | U8 | OP2 | 2.96 | Arg142 | NH2 |
| 25 | U8 | OP2 | 2.84 | Arg142 | NH1 |

**Table S5. Details on atoms of the interacting residues involved in protein-protein interactions of two distant subunits ($N_{n+1}$ and $N_{n-1}$).**
The analysis was done using the PDBePisa tool [2,3]. A salt bridge is highlighted in green; the rest represents H-bonds.

| Number of the contact | $N_{n+1}$ subunit | | Distance (Å) | $N_{n-1}$ subunit | |
|---|---|---|---|---|---|
| | Residue | Atom | | Residue | Atom |
| 1 | Leu347 | O | 3.14 | Tyr2 | N |
| 2 | Glu355 | OE1 | 2.82 | Tyr2 | OH |
| 3 | Ala345 | O | 3.64 | Cys3 | SG |
| 4 | Ala345 | O | 2.86 | Thr4 | N |
| 5 | Lys344 | O | 2.73 | Thr4 | OG1 |
| 6 | Leu347 | N | 2.77 | Tyr2 | O |
| 7 | Ala345 | O | 3.36 | Thr4 | OG1 |
| 8 | Glu355 | OE2 | 2.93 | Lys7 | NZ |

**Table S6. Details on the atoms of the interacting residues involved in protein-protein interactions of two adjacent subunits.**

The analysis was done using the PDBePisa tool [2,3]. A salt bridge is highlighted in green; the rest represents H-bonds.

| Number of the contact | $N_{n+1}$ subunit | | Distance (Å) | N subunit | |
|---|---|---|---|---|---|
| | Residue | Atom | | Residue | Atom |
| 1 | Asp234 | OD2 | 3.27 | His320 | NE2 |
| 2 | Asp234 | O | 2.87 | Asn321 | N |
| 3 | Leu306 | O | 3.6 | Leu323 | NE2 |
| 4 | Thr307 | OG1 | 3.32 | Tyr426 | OH |
| 5 | Arg219 | NH2 | 3.58 | Asp17 | OD1 |
| 6 | Arg219 | NE | 3.33 | Asp17 | OD2 |
| 7 | Lys344 | NZ | 2.88 | Asp245 | O |
| 8 | Lys344 | NZ | 3.05 | Asp245 | OD1 |
| 9 | Lys348 | NZ | 2.9 | Asp245 | OD2 |
| 10 | Ala345 | N | 3.53 | Ile247 | O |
| 11 | Thr343 | N | 2.82 | Phe248 | O |
| 12 | Glu309 | N | 3.24 | Glu319 | OE1 |
| 13 | Ala341 | N | 2.9 | Asn324 | OD1 |
| 14 | Lys348 | NZ | 2.95 | Asp385 | OD1 |
| 15 | Ala340 | N | 3.19 | Phe398 | O |
