## Supplementary Data S1 for "Cryo-electron microscopy structure of the bovine ephemeral fever virus RNA-nucleoprotein assembly"

**Data S1. Phylogenetic tree and multiple sequence alignment based on nucleoprotein sequences of *Mononegavirales* species.**

These include uncharacterised Ephemerovirus genus species (from Berrimah to Yata viruses), VSV, RABV (all from Rhabdoviridae family), and more distant species from other families within the Mononegavirales order, such as Ebola virus (Filoviridae family), Nipah and Measles viruses (Paramyxoviridae family), and Human Respiratory Syncytial virus (Pneumiviridae family).

**Data S1.1. Phylogenetic tree based on nucleoprotein sequences of selected *Mononegavirales* species.**

MOR (Mononegavirales outside Rhabdoviridae) selected as the tree root. The values above the nodes of the branches depict the confidence value corresponding to 100 transfer bootstrap estimates. The scale bar denotes amino acid substitutions/site.

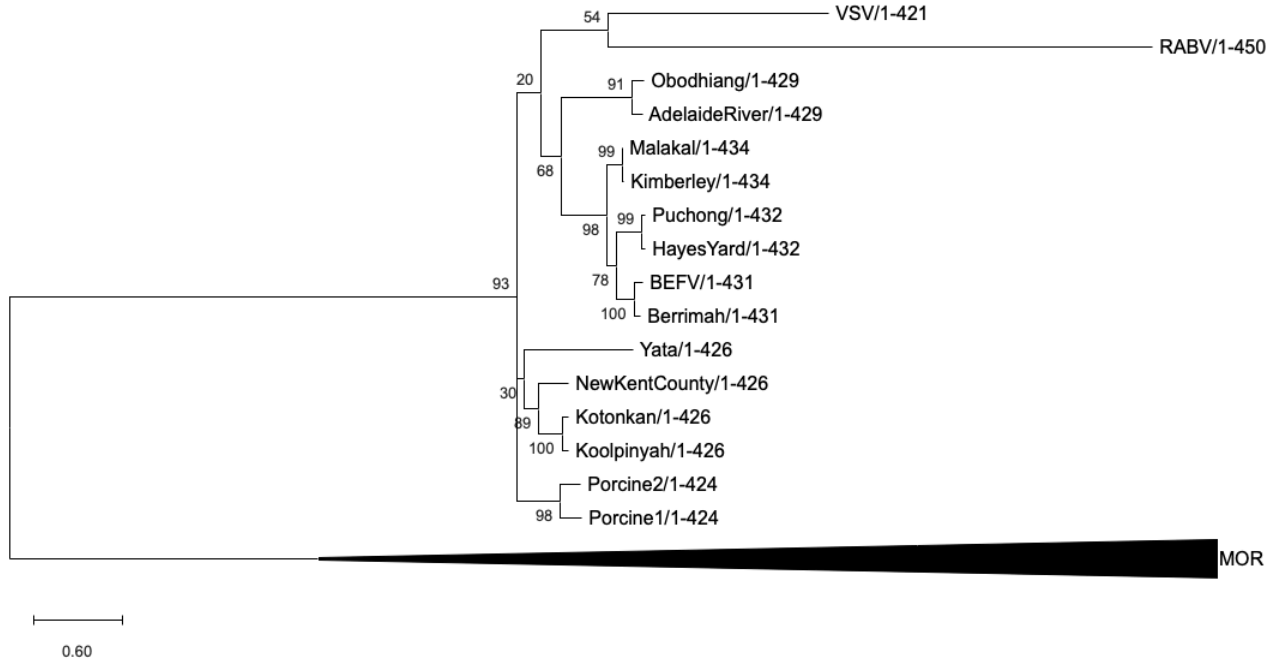


**Data S1.2. Visualisation of multiple sequence alignment of BEFV and selected *Mononegavirales* species.**

Secondary structure of BEFV N is assigned to the sequence shown above the BEFV N sequence. Identical residues are shown in red boxes. Group-similarity residues are shown as red letters in blue boxes on a white background.

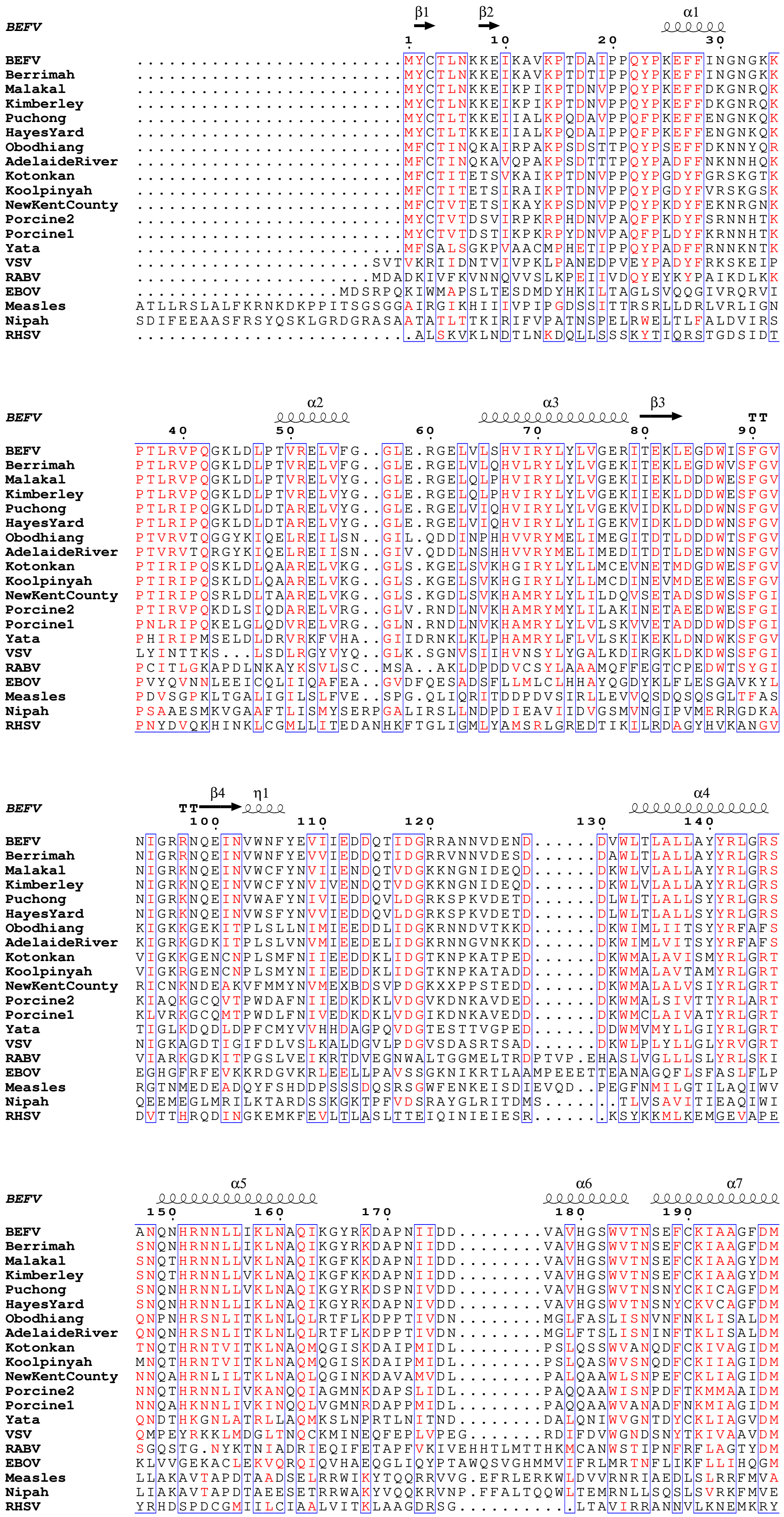
BEFV
β1 β2 α1
1 10 20 30
BEFV .........................MYCTLNKKEIKAVKPTDAIPPQYPKEFFINGNGKK
Berrimah .........................MYCTLNKKEIKAVKPTDTIPPQYPKEFFINGNGKK
Malakal .........................MYCTLNKKEIKPIKPTDNVPPQYPKEFFDKGNRQK
Kimberley .........................MYCTLNKKEIKPIKPTDNVPPQYPKEFFDKGNRQK
Puchong .........................MYCTLTKKEIIALKPQDAVPPQFPKEFFENGNKQK
HayesYard .........................MYCTLTKKEIIALKPQDAIPPQFPKEFFENGNKQK
Obodhiang .........................MFCTINQKAIRPAKPSDSTTPQYPSEFFDKNNYQR
AdelaideRiver .........................MFCTINQKAVQPAKPSDTTTPQYPADFFNKNNHQK
Kotonkan .........................MFCTITETSVKAIKPTDNVPPQYPGDYFGRSKGTK
Koolpinyah .........................MFCTITETSIRAIKPTDNVPPQYPGDYFVRSKGSK
NewKentCounty .........................MFCTVTETSIKAYKPSDNVPPQYPKDYFEKNRGNK
Porcine2 .........................MYCTVTDSVIRPKRPHDNVPAQFPKDYFSRNNHTK
Porcine1 .........................MYCTVTDSTIKPKRPYDNVPAQFPLDYFKRNNHTK
Yata .........................MFSALSGKPVAACMPHETIPPQYPADFFRNNKNTK
VSV ......................SVTVKRIIDNTVIVPKLPANEDPVEYPADYFRKSKEIP
RABV ......................MDADKIVFKVNNQVVSLKPEIIVDQYEYKYPAIKDLKK
EBOV ...................MDSRPQKIWMAPSLTESDMDYHKILTAGLSVQQGIVRQRVI
Measles ATLLRSLALFKRNKDKPPITSGSGGAIRGIKHIIIVPIPGDSSITTRSRLLDRLVRLIGN
Nipah SDIFEEAASFRSYQSKLGRDGRASAATATLTTKIRIFVPATNSPELRWELTLFALDVIRS
RHSV ..........................ALSKVKLNDTLNKDQLLSSSKYTIQRSTGDSIDT
BEFV
α2 α3 β3 TT
40 50 60 70 80 90
BEFV PTLRVPQGKLDLPTVRELVFG..GLE.RGELVLSHVIRYLYLVGERITEKLEGDWISFGV
Berrimah PTLRVPQGKLDLPTVRELVFG..GLE.RGELVLQHVLRYLYLVGEKITEKLEGDWVSFGV
Malakal PTLRVPQGKLDLPTVRELVYG..GLE.RGELQLPHVIRYLYLVGEKIIEKLDDDWESFGV
Kimberley PTLRVPQGKLDLPTVRELVYG..GLE.RGELQLPHVIRYLYLVGEKIIEKLDDDWESFGV
Puchong PTLRIPQGKLDLDTARELVYG..GLE.RGELVIQHVIRYLYLVGEKVIDKLDDDWNSFGV
HayesYard PTLRIPQGKLDLDTARELVYG..GLE.RGELVIQHVIRYLYLIGEKVIDKLDDDWNSFGV
Obodhiang PTVRVTQGGYKIQELREILSN..GIL.QDDINPHHVVRYMELIMEGITDTLDDDWTSFGV
AdelaideRiver PTVRVTQRGYKIQELREIISN..GIV.QDDLNSHHVVRYMELIMEDITDTLDEDWNSFGV
Kotonkan PTIRIPQSKLDLQAARELVKG..GLS.KGELSVKHGIRYLYLLMCEVNETMDGDWESFGV
Koolpinyah PTIRIPQSKLDLQAARELVKG..GLS.KGELSVKHGIRYLYLLMCDINEVMDEEWESFGV
NewKentCounty PTIRIPQSRLDLTAARELVKG..GLS.KGDLSVKHAMRYLYLILDQVSETADSDWSSFGI
Porcine2 PTIRVPQKDLSIQDARELVRG..GLV.RNDLNVKHAMRYMYLILAKINETAEEDWESFGI
Porcine1 PNLRIPQKELGLQDVRELVRG..GLN.RNDLNVKHAMRYLYLVLSKVVETADDDWDSFGI
Yata PHIRIPMSELDLDRVRKFVHA..GIIDRNKLKLPHAMRYLFLVLSKIKEKLDNDWKSFGV
VSV LYINTTKS...LSDLRGYVYQ..GLK.SGNVSIIHVNSYLYGALKDIRGKLDKDWSSFGI
RABV PCITLGKAPDLNKAYKSVLSC..MSA..AKLDPDDVCSYLAAAMQFFEGTCPEDWTSYGI
EBOV PVYQVNNLEEICQLIIQAFEA..GVDFQESADSFLLMLCLHHAYQGDYKLFLESGAVKYL
Measles PDVSGPKLTGALIGILSLFVE..SPG.QLIQRITDDPDVSIRLLEVVQSDQSQSGLTFAS
Nipah PSAAESMKVGAAFTLISMYSERPGALIRSLLNDPDIEAVIIDVGSMVNGIPVMERRGDKA
RHSV PNYDVQKHINKLCGMLLITEDANHKFTGLIGMLYAMSRLGREDTIKILRDAGYHVKANGV
BEFV
TT β4 η1 α4
100 110 120 130 140
BEFV NIGRRNQEINVWNFYEVIIEDDQTIDGRRANNVDEND......DVWLTLALLAYYRLGRS
Berrimah NIGRRNQEINVWNFYEVVIEDDQTIDGRRVNNVDESD......DAWLTLALLAYYRLGRS
Malakal NIGRKNQEINVWCFYNVIIENDQTVDGKKNGNIDEQD......DKWLVLALLAYYRLGRS
Kimberley NIGRKNQEINVWCFYNVIVENDQTVDGKKNGNIDEQD......DKWLVLALLAYYRLGRS
Puchong NIGRKNQEINVWAFYNIVIEDDQVLDGRKSPKVDETD......DLWLTLALLSYYRLGRS
HayesYard NIGRKNQEINVWSFYNVVIEDDQVLDGRKSPKVDETD......DLWLTLALLSYYRLGRS
Obodhiang KIGKKGEKITPLSLLNIMIEEDDLIDGKRNNDVTKKD......DKWIMLIITSYYRFAFS
AdelaideRiver KIGRKGDKITPLSLVNVMIEEDELIDGKRNNGVNKKD......DKWIMLVITSYYRFAFS
Kotonkan VIGKKGENCNPLSMFNIIEEDDKLIDGTKNPKATPED......DKWMALAVISMYRLGRT
Koolpinyah VIGKRGENCNPLSMYNIIEEDDKLIDGTKNPKATADD......DKWMALAVTAMYRLGRT
NewKentCounty RICNKNDEAKVFMMYNVMEXBDSVPDGKXXPPSTEDD......DKWMALALVSIYRLGRT
Porcine2 KIAQKGCQVTPWDAFNIIEDKDKLVDGVKDNKAVDED......DKWMALSIVTTYRLART
Porcine1 KLVRKGCQMTPWDLFNIVEDKDKLVDGIKDNKAVEDD......DKWMCLAIVATYRLGRT
Yata TIGLKDQDLDPFCMYVVHHDAGPQVDGTESTTVGPED......DDWMVMYLLGIYRLGRT
VSV NIGKAGDTIGIFDLVSLKALDGVLPDGVSDASRTSAD......DKWLPLYLLGLYRVGRT
RABV VIARKGDKITPGSLVEIKRTDVEGNWALTGGMELTRDPTVP.EHASLVGLLLSLYRLSKI
EBOV EGHGFRFEVKKRDGVKRLEELLPAVSSGKNIKRTLAAMPEEETTEANAGQFLSFASLFLP
Measles RGTNMEDEADQYFSHDDPSSSDQSRSGWFENKEISDIEVQD..PEGFNMILGTILAQIWV
Nipah QEEMEGLMRILKTARDSSKGKTPFVDSRAYGLRITDMS.......TLVSAVITIEAQIWI
RHSV DVTTHRQDINGKEMKFEVLTLASLTTEIQINIEIESR.......KSYKKMLKEMGEVAPE
BEFV
α5 α6 α7
150 160 170 180 190
BEFV ANQNHRNNLLIKLNAQIKGYRKDAPNIIDD........VAVHGSWVTNSEFCKIAAGFDM
Berrimah SNQNHRNNLLIKLNAQIKGYRKDAPNIIDD........VAVHGSWVTNSEFCKIAAGFDM
Malakal SNQTHRNNLLVKLNAQIKGFKKDAPNIIDD........VAVHGSWVTNSEFCKIAAGYDM
Kimberley SNQTHRNNLLVKLNAQIKGFKKDAPNIIDD........VAVHGSWVTNSEFCKIAAGYDM
Puchong SNQNHRNNLLVKLNAQIKGYRKDSPNIVDD........VAVHGSWVTNSNYCKICAGFDM
HayesYard SNQNHRNNLLIKLNAQIKGYRKDAPNIVDD........VAVHGSWVTNSNYCKVCAGFDM
Obodhiang QNPNHRSNLITKLNLQLRTFLKDPPTIVDN........MGLFASLISNVNFNKLISALDM
AdelaideRiver QNQNHRSNLITKLNLQLRTFLKDPPTIVDN........MGLFTSLISNINFTKLISALDM
Kotonkan TNQTHRNTVITKLNAQMQGISKDAIPMIDL........PSLQSSWVANQDFCKIVAGIDM
Koolpinyah MNQTHRNTVITKLNAQMQGISKDAIPMIDL........PSLQASWVSNQDFCKIIAGIDM
NewKentCounty NNQAHRNLILTKLNAQLQGINKDAVAMVDL........PALQAAWLSNPEFCKLIAGIDM
Porcine2 NNQTHRNNLIVKANQQIAGMNKDAPSLIDL........PAQQAAWISNPDFTKMMAAIDM
Porcine1 NNQAHKNNLIVKINQQLVGMNRDAPPMIDL........PAQQAAWVANADFNKMIAGIDM
Yata QNDTHKGNLATRLLAQMKSLNPRTLNITND........DALQNIWVGNTDYCKLIAGVDM
VSV QMPEYRKKLMDGLTNQCKMINEQFEPLVPEG.......RDIFDVWGNDSNYTKIVAAVDM
RABV SGQSTG.NYKTNIADRIEQIFETAPFVKIVEHHTLMTTHKMCANWSTIPNFRFLAGTYDM
EBOV KLVVGEKACLEKVQRQIQVHAEQGLIQYPTAWQSVGHMMVIFRLMRTNFLIKFLLIHQGM
Measles LLAKAVTAPDTAADSELRRWIKYTQQRRVVG.EFRLERKWLDVVRNRIAEDLSLRRFMVA
Nipah LIAKAVTAPDTAEESETRRWAKYVQQKRVNP.FFALTQQWLTEMRNLLSQSLSVRKFMVE
RHSV YRHDSPDCGMIILCIAALVITKLAAGDRSG...........LTAVIRRANNVLKNEMKRY

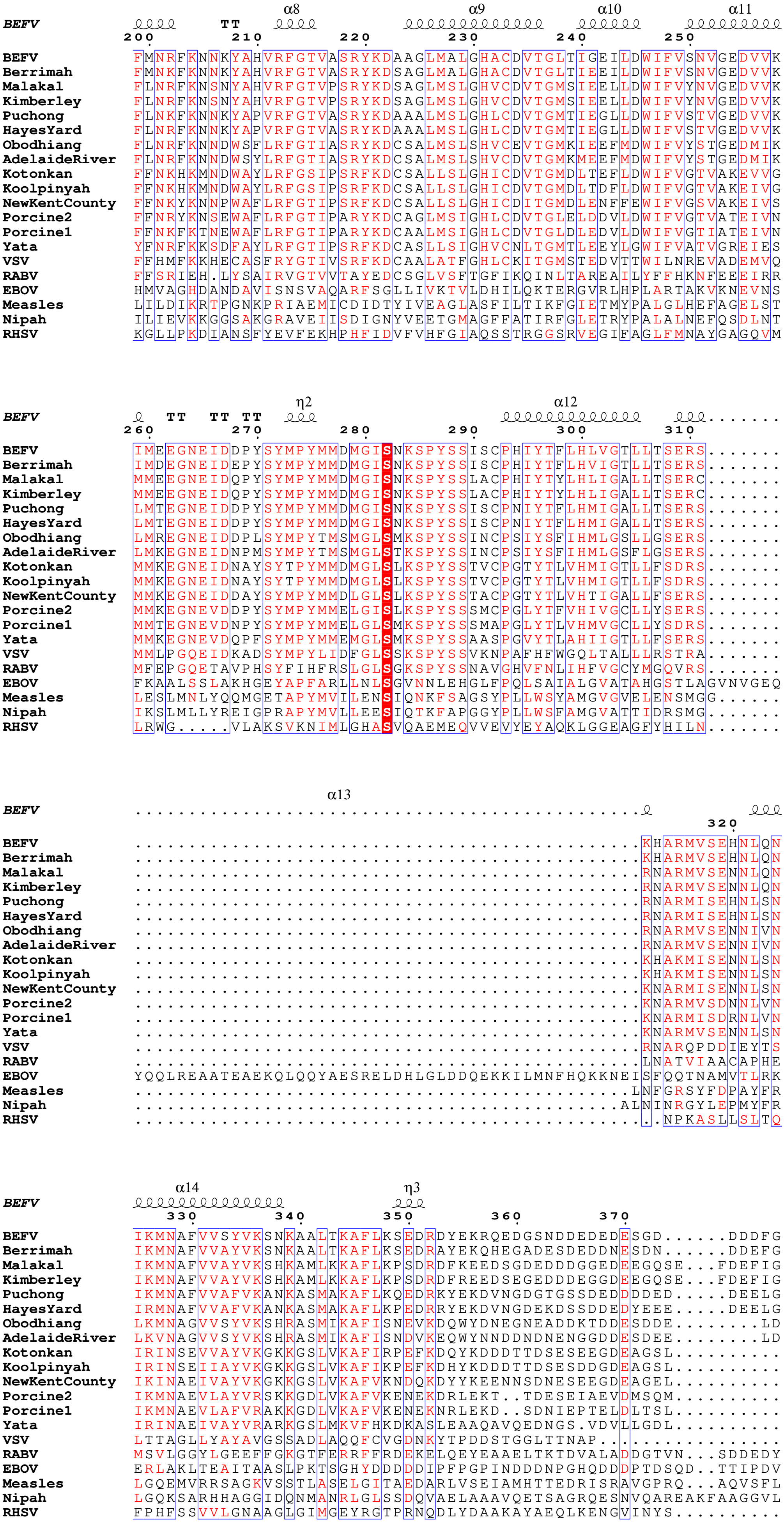
BEFV
α8 α9 α10 α11
TT
200 210 220 230 240 250
BEFV FMNRFKNNKYAHVRFGTVASRYKDAAGLMALGHACDVTGLTIGEILDWIFVSNVGEDVVK
Berrimah FMNKFKNNKYAHVRFGTVASRYKDSAGLMALGHACDVTGLTIEEILDWIFVSNVGEDVVK
Malakal FLNRFKNSNYAHVRFGTVPSRYKDSAGLMSLGHVCDVTGMSIEELLDWIFVYNVGEDVVK
Kimberley FLNRFKNSNYAHVRFGTVPSRYKDSAGLMSLGHVCDVTGMSIEELLDWIFVYNVGEDVVK
Puchong FLNKFKNNKYAPVRFGTVASRYKDAAALMSLGHLCDVTGMTIEGLLDWIFVSTVGEDVVK
HayesYard FLNRFKNNKYAPVRFGTVASRYKDAAALMSLGHLCDVTGMTIEGLLDWIFVSTVGEDVVK
Obodhiang FLNRFKNNDWSFLRFGTIASRYKDCSALMSLSHVCEVTGMKIEEFMDWIFVYSTGEDMIK
AdelaideRiver FLNRFKNNDWSYLRFGTIASRYKDCSALMSLSHVCDVTGMKMEEFMDWIFVYSTGEDMIK
Kotonkan FFNKHKMNDWAYLRFGSIPSRFKDCSALLSLGHICDVTGMDLTEFLDWIFVGTVAKEVVG
Koolpinyah FFNKHKMNDWAYLRFGSIPSRFKDCSALLSLGHICDVTGMDLTDFLDWIFVGTVAKEIVG
NewKentCounty FFNKYKNNPWAFLRFGTIPSRFKDCSALLSLGHICDITGMDLENFFEWIFVGSVAKEIVS
Porcine2 FFNRYKNSEWAFLRFGTIPARYKDCAGLMSIGHLCDVTGLELDDVLDWIFVGTVATEIVN
Porcine1 FFNKFKTNEWAFLRFGTIPARYKDCAALMSIGHLCDVTGLDLEDVLDWIFVGTIATEIVN
Yata YFNRFKKSDFAYLRFGTIPSRFKDCASLLSIGHVCNLTGMTLEEYLGWIFVATVGREIES
VSV FFHMFKKHECASFRYGTIVSRFKDCAALATFGHLCKITGMSTEDVTTWILNREVADEMVQ
RABV FFSRIEH.LYSAIRVGTVVTAYEDCSGLVSFTGFIKQINLTAREAILYFFHKNFEEEIRR
EBOV HMVAGHDANDAVISNSVAQARFSGLLIVKTVLDHILQKTERGVRLHPLARTAKVKNEVNS
Measles LILDIKRTPGNKPRIAEMICDIDTYIVEAGLASFILTIKFGIETMYPALGLHEFAGELST
Nipah ILIEVKKGGSAKGRAVEIISDIGNYVEETGMAGFFATIRFGLETRYPALALNEFQSDLNT
RHSV KGLLPKDIANSFYEVFEKHPHFIDVFVHFGIAQSSTRGGSRVEGIFAGLFMNAYGAGQVM

BEFV
η2 α12
TT TT TT
260 270 280 290 300 310
BEFV IMEEGNEIDDPYSYMPYMMDMGISNKSPYSSISCPHIYTFLHLVGTLLTSERS.......
Berrimah IMDEGNEIDEPYSYMPYMMDMGISNKSPYSSISCPHIYTFLHVIGTLLTSERS.......
Malakal MMEEGNEIDQPYSYMPYMMDMGISNKSPYSSLACPHIYTYLHLIGALLTSERC.......
Kimberley MMEEGNEIDQPYSYMPYMMDMGISNKSPYSSLACPHIYTYLHLIGALLTSERC.......
Puchong LMTEGNEIDDPYSYMPYMMDMGISNKSPYSSISCPNIYTFLHMIGALLTSERS.......
HayesYard LMTEGNEIDDPYSYMPYMMDMGISNKSPYSSISCPNIYTFLHMIGALLTSERS.......
Obodhiang LMREGNEIDDPLSYMPYTMSMGLSMKSPYSSINCPSIYSFIHMLGSLLGSERS.......
AdelaideRiver LMKEGNEIDNPMSYMPYTMSMGLSTKSPYSSINCPSIYSFIHMLGSFLGSERS.......
Kotonkan MMKEGNEIDNAYSYTPYMMDMGLSLKSPYSSTVCPGTYTLVHMIGTLLFSDRS.......
Koolpinyah MMKEGNEIDNAYSYTPYMMDMGLSLKSPYSSTVCPGTYTLVHMIGTLLFSDRS.......
NewKentCounty MMKEGNEIDDAYSYMPYMMDLGLSLKSPYSSTACPGTYTLVHTIGALLFSERS.......
Porcine2 MMKEGNEVDDPYSYMPYMMELGISLKSPYSSSMCPGLYTFVHIVGCLLYSERS.......
Porcine1 MMTEGNEVDNPYSYMPYMMELGLSMKSPYSSSMAPGLYTLVHMVGCLLYSDRS.......
Yata MMKEGNEVDQPFSYMPYMMEMGLSMKSPYSSAASPGVYTLAHIIGTLLFSERS.......
VSV MMLPGQEIDKADSYMPYLIDFGLSSKSPYSSVKNPAFHFWGQLTALLLRSTRA.......
RABV MFEPGQETAVPHSYFIHFRSLGLSGKSPYSSNAVGHVFNLIHFVGCYMGQVRS.......
EBOV FKAALSSLAKHGEYAPFARLLNLSGVNNLEHGLFPQLSAIALGVATAHGSTLAGVNVGEQ
Measles LESLMNLYQQMGETAPYMVILENSIQNKFSAGSYPLLWSYAMGVGVELENSMGG......
Nipah IKSLMLLYREIGPRAPYMVLLEESIQTKFAPGGYPLLWSFAMGVATTIDRSMG.......
RHSV LRWG.....VLAKSVKNIMLGHASVQAEMEQVVEVYEYAQKLGGEAGFYHILN.......

BEFV
α13
320
BEFV ...............................................KHARMVSEHNLQN
Berrimah ...............................................KHARMVSEHNLQN
Malakal ...............................................RNARMVSENNLQN
Kimberley ...............................................RNARMVSENNLQN
Puchong ...............................................RNARMISEHNLSN
HayesYard ...............................................RNARMISEHNLSN
Obodhiang ...............................................RNARMVSENNIVN
AdelaideRiver ...............................................RNARMVSENNIVN
Kotonkan ...............................................KHAKMISENNLSN
Koolpinyah ...............................................KHAKMISENNLSN
NewKentCounty ...............................................KNARMISENNLSN
Porcine2 ...............................................KNARMVSDNNLVN
Porcine1 ...............................................KNARMISDRNLVN
Yata ...............................................KNARMVSENNLSN
VSV ...............................................RNARQPDDIEYTS
RABV ...............................................LNATVIAACAPHE
EBOV YQQLREAATEAEKQLQQYAESRELDHLGLDDQEKKILMNFHQKKNEISFQQTNAMVTLRK
Measles ..............................................LNFGRSYFDPAYFR
Nipah .............................................ALNINRGYLEPMYFR
RHSV .................................................NPKASLLSLTQ

BEFV
α14 η3
330 340 350 360 370
BEFV IKMNAFVVSYVKSNKAALTKAFLKSEDRDYEKRQEDGSNDDEDEDESGD......DDDFG
Berrimah IKMNAFVVAYVKSNKAALTKAFLKSEDRAYEKQHEGADESDEDDNESDN......DDEFG
Malakal IKMNAFVVAYVKSHKAMLKKAFLKPSDRDFKEEDSGDEDDDGGEDEEGQSE...FDEFIG
Kimberley IKMNAFVVAYVKSHKAMLKKAFLKPSDRDFREEDSEGEDDDEGGDEEGQSE...FDEFIG
Puchong IKMNAFVVAFVKANKASMAKAFLKQEDRKYEKDVNGDGTGSSDEDDDDED.....DEELG
HayesYard IRMNAFVVAFVKANKASMAKAFLKPEDRRYEKDVNGDEKDSSDDEDYEEE.....DEELG
Obodhiang LKMNAGVVSYVKSHRASMIKAFISNEVKDQWYDNEGNEADDKTDDESDDE........LD
AdelaideRiver LKVNAGVVSYVKSHRASMIKAFISNDVKEQWYNNDDNDNENGGDDESDEE........LD
Kotonkan IRINSEVVAYVKGKKGSLVKAFIRPEFKDQYKDDDTTDSESEEGDEAGSL..........
Koolpinyah IRINSEIIAYVKGKKGSLVKAFIKPEFKDHYKDDDTTDSESDDGDEAGSL..........
NewKentCounty IKINAEIVAYVKGKKGSLVKAFVKNDQKDYYKEENNSDNESEEGDEAGEL..........
Porcine2 IKMNAEVLAYVRSKKGDLVKAFVKENEKDRLEKT.TDESEIAEVDMSQM...........
Porcine1 IKMNAEVLAFVRAKKGDLVKAFVKENEKNRLEKD..SDNIEPTELDLTSL..........
Yata IRINAEIVAYVRARKGSLMKVFHKDKASLEAAQAVQEDNGS.VDVLLGDL..........
VSV LTTAGLLYAYAVGSSADLAQQFCVGDNKYTPDDSTGGLTTNAP.................
RABV MSVLGGYLGEEFFGKGTFERRFFRDEKELQEYEAAELTKTDVALADDGTVN...SDDEDY
EBOV ERLAKLTEAITAASLPKTSGHYDDDDDIPFPGPINDDDNPGHQDDDPTDSQD..TTIPDV
Measles LGQEMVRRSAGKVSSTLASELGITAEDARLVSEIAMHTTEDRISRAVGPRQ...AQVSFL
Nipah LGQKSARHHAGGIDQNMANRLGLSSDQVAELAAAVQETSAGRQESNVQAREAKFAAGGVL
RHSV FPHFSSVVLGNAAGLGIMGEYRGTPRNQDLYDAAKAYAEQLKENGVINYS..........

```
                                  α15                                  α16
BEFV                        ℓℓℓℓℓℓℓℓℓℓ      ℓℓℓℓℓℓℓℓℓℓ...........................
                  380       390       400       410
BEFV              AMPKSSDPMEWFIFLESNHFILPEKVTEFCIRE...........................
Berrimah          AMPKSSDPMEWFIYLESNHFVLPDKVTEFCMRE...........................
Malakal           DMPKSSNPMEWYIYLQSNHFALPDKVVDFCLKE...........................
Kimberley         DMPKSSNPMEWYIYLQSNHFALPDKVVDFCLKE...........................
Puchong           DMPKSADPMEWFVYLQAQHFTLPDKVNEFGQRE...........................
HayesYard         DMPKSADPMEWFVYLQAQHFTLPDKVNDFGQKE...........................
Obodhiang         GMPKGDNPVEWFMFLESRHFELPDEIKSFMNRE...........................
AdelaideRiver     EMPKGDNPVEWFMYLESRHFELPEEIKNFMNRE...........................
Kotonkan          ..PRTDDPMEWFAYLEYNHFDLPDVIKEHTRSE...........................
Koolpinyah        ..PKSDDPMEWFAFLEYNHFDLPDVIKEHTRLE...........................
NewKentCounty     ..PSSDDAMEWFAYLEXXHFDLPDIIKEHTRIE...........................
Porcine2          ..PTSNDPMEWFAYLEFMDFNLPEEIKHHTKSE...........................
Porcine1          ..PSSHDPMDWFAYLESVDFNLPDEIKLHTKSE...........................
Yata              ..PAGNDPDEWFTTLEINQFELPHEIRTHTISE...........................
VSV               ..PQGRDVVEWLGWFEDQNRKPTPDMMQYAKRA...........................
RABV              FSGETRSPEAVYTRIIMNGGRLKRSHIRRYVSV...........................
EBOV              VVDPDDGSYGEYQSYSENGMNAPDDLVLFDLDEDDEDTKPVPNRSTKGGQQKNSQKGQHI
Measles           HGDQSENELPGLGGKEDRRVKQGRGEARESYRETG.........................
Nipah             IGGSDQDIDEGEEPIEQSGRQSVTFKREMSISSLAN........................
RHSV              ...VLDLTAEELEAIKHQLNPKDNDVEL................................

                                   α17
BEFV              ......ℓℓ       ℓℓℓℓℓℓ
                            420       430
BEFV              ......CKKIQNARPNTIGKYLASIV..................................
Berrimah          ......CKKIQNARPNTIGKYLSSIV..................................
Malakal           ......AKKIQNARPGTVGKYLSTIA..................................
Kimberley         ......AKKIQNARPGTVGKYLSTIA..................................
Puchong           ......CKKIQNARPGTIGKYLSTIG..................................
HayesYard         ......CKKIQNARPGTIGKYLSTIG..................................
Obodhiang         ......AKKITNPRSGTIGKFVSLMN..................................
AdelaideRiver     ......ARKITNPRVGTIGKFVSTMN..................................
Kotonkan          ......SRKIQNTRAGTIGNHVVTTFN.................................
Koolpinyah        ......SRKIQNTRAGTIGNHVVTTFN.................................
NewKentCounty     ......SKKLTGTRAGTIGYHIVNTFN.................................
Porcine2          ......SKKITNTRAGTIGNHVSTTYC.................................
Porcine1          ......SRKITNTRAGTIGHHIATTFI.................................
Yata              ......SRKIGNTRAHTIGHHVATTFV.................................
VSV               ......VMSLQGLREKTIGKYAKSEFDK................................
RABV              ......SSNHQARPNSFAEFLNKTYSSDS...............................
EBOV              EGRQTQSRPIQNVPGPHRTIHHASAPLTDNDRRNEPSGSTSPRMLTPINEEADPLDDADD
Measles           ......SSRASDARAAHPPTSMPLDIDTASESGQDPQDSRRSADALLRLQAMAGILEEQG
Nipah             ....SVPSSSVSTSGGTRLTNSLLNLRSRLAAKAAKEAASSNATDDPAISNRTQGESEKK
RHSV              ............................................................

BEFV

BEFV              ............................................................
Berrimah          ............................................................
Malakal           ............................................................
Kimberley         ............................................................
Puchong           ............................................................
HayesYard         ............................................................
Obodhiang         ............................................................
AdelaideRiver     ............................................................
Kotonkan          ............................................................
Koolpinyah        ............................................................
NewKentCounty     ............................................................
Porcine2          ............................................................
Porcine1          ............................................................
Yata              ............................................................
VSV               ............................................................
RABV              ............................................................
EBOV              ETSSLPPLESDDEEQDRDGTSNRTPTVAPPAPVYRDHSEKKELPQDEQQDQDHTQEARNQ
Measles           SDTDTPRVYNDRDLLD............................................
Nipah             NNQDLKPAQNDLDFVRADV.........................................
RHSV              ............................................................

BEFV

BEFV              ............................................................
Berrimah          ............................................................
Malakal           ............................................................
Kimberley         ............................................................
Puchong           ............................................................
HayesYard         ............................................................
Obodhiang         ............................................................
AdelaideRiver     ............................................................
Kotonkan          ............................................................
Koolpinyah        ............................................................
NewKentCounty     ............................................................
Porcine2          ............................................................
Porcine1          ............................................................
Yata              ............................................................
VSV               ............................................................
RABV              ............................................................
EBOV              DSDNTQSEHSFEEMYRHILRSQGPFDAVLYYHMMKDEPVVFSTSDGKEYTYPDSLEEEYP
Measles           ............................................................
Nipah             ............................................................
RHSV              ............................................................
```

***BEFV***

```
BEFV            . . . . . . . . . . . . . . . . . . . . . . . . . . . . . . . . . . . . . . . . . .
Berrimah        . . . . . . . . . . . . . . . . . . . . . . . . . . . . . . . . . . . . . . . . . .
Malakal         . . . . . . . . . . . . . . . . . . . . . . . . . . . . . . . . . . . . . . . . . .
Kimberley       . . . . . . . . . . . . . . . . . . . . . . . . . . . . . . . . . . . . . . . . . .
Puchong         . . . . . . . . . . . . . . . . . . . . . . . . . . . . . . . . . . . . . . . . . .
HayesYard       . . . . . . . . . . . . . . . . . . . . . . . . . . . . . . . . . . . . . . . . . .
Obodhiang       . . . . . . . . . . . . . . . . . . . . . . . . . . . . . . . . . . . . . . . . . .
AdelaideRiver   . . . . . . . . . . . . . . . . . . . . . . . . . . . . . . . . . . . . . . . . . .
Kotonkan        . . . . . . . . . . . . . . . . . . . . . . . . . . . . . . . . . . . . . . . . . .
Koolpinyah      . . . . . . . . . . . . . . . . . . . . . . . . . . . . . . . . . . . . . . . . . .
NewKentCounty   . . . . . . . . . . . . . . . . . . . . . . . . . . . . . . . . . . . . . . . . . .
Porcine2        . . . . . . . . . . . . . . . . . . . . . . . . . . . . . . . . . . . . . . . . . .
Porcine1        . . . . . . . . . . . . . . . . . . . . . . . . . . . . . . . . . . . . . . . . . .
Yata            . . . . . . . . . . . . . . . . . . . . . . . . . . . . . . . . . . . . . . . . . .
VSV             . . . . . . . . . . . . . . . . . . . . . . . . . . . . . . . . . . . . . . . . . .
RABV            . . . . . . . . . . . . . . . . . . . . . . . . . . . . . . . . . . . . . . . . . .
EBOV            P W L T E K E A M N E E N R F V T L D G Q Q F Y W P V M N H K N K F M A I L Q H H Q
Measles         . . . . . . . . . . . . . . . . . . . . . . . . . . . . . . . . . . . . . . . . . .
Nipah           . . . . . . . . . . . . . . . . . . . . . . . . . . . . . . . . . . . . . . . . . .
RHSV            . . . . . . . . . . . . . . . . . . . . . . . . . . . . . . . . . . . . . . . . . .
```

**Data S1.3. Raw multiple sequence alignment data of BEFV and selected *Mononegavirales* species.**

>BEFV/1-431
------------------------MYCTLNKKEIKAVKPTDAIPPQYPKEFFINGNGKKPTLRVPQGKLDLPTV
RELVFG--GLE-RGELVLSHVIRYLYLVGERITEKLEGDWISFGVNIGRRNQEINVWNFYEVIIE
DDQTIDGRRANNVDEND------DVWLTLALLAYYRLGRSANQNHRNNLLIKLNAQIKGYRKDA
PNIIDD--------VAVHGSWVTNSEFCKIAAGFDMFMNRFKNNKYAHVRFGTVASRYKDAAGLM
ALGHACDVTGLTIGEILDWIFVSNVGEDVVKIMEEGNEIDDPYSYMPYMMDMGISNKSPYSS
ISCPHIYTFLHLVGTLLTSERS------------------------------------------------------KHARMVSEHNLQNI
KMNAFVVSYVKSNKAALTKAFLKSEDRDYEKRQEDGSNDDEDEDESGD------DDDFGAMP
KSSDPMEWFIFLESNHFILPEKVTEFCIRE---------------------------------CKKIQNARPNTIGKYLA
SIV----------------------------------------------------------------------------------------------------------------------
------------------------------------------------------------------------------

>Berrimah/1-431
------------------------MYCTLNKKEIKAVKPTDTIPPQYPKEFFINGNGKKPTLRVPQGKLDLPTV
RELVFG--GLE-RGELVLQHVLRYLYLVGEKITEKLEGDWVSFGVNIGRRNQEINVWNFYEVVI
EDDQTIDGRRVNNVDESD------DAWLTLALLAYYRLGRSSNQNHRNNLLIKLNAQIKGYRKD
APNIIDD--------VAVHGSWVTNSEFCKIAAGFDMFMNKFKNNKYAHVRFGTVASRYKDSAGL
MALGHACDVTGLTIEEILDWIFVSNVGEDVVKIMDEGNEIDEPYSYMPYMMDMGISNKSPYS
SISCPHIYTFLHVIGTLLTSERS------------------------------------------------------KHARMVSEHNLQN
IKMNAFVVAYVKSNKAALTKAFLKSEDRAYEKQHEGADESDEDDNESDN------DDEFGAMPK
SSDPMEWFIYLESNHFVLPDKVTEFCMRE---------------------------------CKKIQNARPNTIGKYL
SSIV--------------------------------------------------------------------------------------------------------------------
--------------------------------------------------------------------------------

>Malakal/1-434
------------------------MYCTLNKKEIKPIKPTDNVPPQYPKEFFDKGNRQKPTLRVPQGKLDLPT
VRELVYG--GLE-RGELQLPHVIRYLYLVGEKIIEKLDDDWESFGVNIGRKNQEINVWCFYNVII
ENDQTVDGKKNGNIDEQD------DKWLVLALLAYYRLGRSSNQTHRNNLLVKLNAQIKGFKKD
APNIIDD--------VAVHGSWVTNSEFCKIAAGYDMFLNRFKNSNYAHVRFGTVPSRYKDSAGL
MSLGHVCDVTGMSIEELLDWIFVYNVGEDVVKMMEEGNEIDQPYSYMPYMMDMGISNKSP
YSSLACPHIYTYLHLIGALLTSERC------------------------------------------------------RNARMVSENN
LQNIKMNAFVVAYVKSHKAMLKKAFLKPSDRDFKEEDSGDEDDDGGEDEEGQSE---FDEFI
GDMPKSSNPMEWYIYLQSNHFALPDKVVDFCLKE---------------------------------AKKIQNARPG
TVGKYLSTIA--------------------------------------------------------------------------------------------------------
--------------------------------------------------------------------------------------------

>Kimberley/1-434
------------------------MYCTLNKKEIKPIKPTDNVPPQYPKEFFDKGNRQKPTLRVPQGKLDLPT
VRELVYG--GLE-RGELQLPHVIRYLYLVGEKIIEKLDDDWESFGVNIGRKNQEINVWCFYNVIV
ENDQTVDGKKNGNIDEQD------DKWLVLALLAYYRLGRSSNQTHRNNLLVKLNAQIKGFKKD
APNIIDD--------VAVHGSWVTNSEFCKIAAGYDMFLNRFKNSNYAHVRFGTVPSRYKDSAGL
MSLGHVCDVTGMSIEELLDWIFVYNVGEDVVKMMEEGNEIDQPYSYMPYMMDMGISNKSP
YSSLACPHIYTYLHLIGALLTSERC------------------------------------------------------RNARMVSENN
LQNIKMNAFVVAYVKSHKAMLKKAFLKPSDRDFREEDSEGEDDDEGGDEEGQSE---FDEFI
GDMPKSSNPMEWYIYLQSNHFALPDKVVDFCLKE---------------------------------AKKIQNARPG
TVGKYLSTIA----------

------------------------------------------------------------------------------------------------------------------------------
----------------------------------------------------------------
>Puchong/1-432
------------------------MYCTLTKKEIIALKPQDAVPPQFPKEFFENGNKQKPTLRIPQGKLDLDTA
RELVYG--GLE-RGELVIQHVIRYLYLVGEKVIDKLDDDWNSFGVNIGRKNQEINVWAFYNIVIE
DDQVLDGRKSPKVDETD------DLWLTLALLSYYRLGRSSNQNHRNNLLVKLNAQIKGYRKDS
PNIVDD--------VAVHGSWVTNSNYCKICAGFDMFLNKFKNNKYAPVRFGTVASRYKDAAALM
SLGHLCDVTGMTIEGLLDWIFVSTVGEDVVKLMTEGNEIDDPYSYMPYMMDMGISNKSPYS
SISCPNIYTFLHMIGALLTSERS-------------------------------------------------------RNARMISEHNLSNI
KMNAFVVAFVKANKASMAKAFLKQEDRKYEKDVNGDGTGSSDEDDDDED-----DEELGDMP
KSADPMEWFVYLQAQHFTLPDKVNEFGQRE---------------------------------CKKIQNARPGTIGK
YLSTIG-------------------------------------------------------------------------------------------------------------------
--------------------------------------------------------------------------------------
>HayesYard/1-432
------------------------MYCTLTKKEIIALKPQDAIPPQFPKEFFENGNKQKPTLRIPQGKLDLDTAR
ELVYG--GLE-RGELVIQHVIRYLYLIGEKVIDKLDDDWNSFGVNIGRKNQEINVWSFYNVVIED
DQVLDGRKSPKVDETD------DLWLTLALLSYYRLGRSSNQNHRNNLLIKLNAQIKGYRKDAP
NIVDD--------VAVHGSWVTNSNYCKVCAGFDMFLNRFKNNKYAPVRFGTVASRYKDAAALM
SLGHLCDVTGMTIEGLLDWIFVSTVGEDVVKLMTEGNEIDDPYSYMPYMMDMGISNKSPYS
SISCPNIYTFLHMIGALLTSERS-------------------------------------------------------RNARMISEHNLSNI
RMNAFVVAFVKANKASMAKAFLKPEDRRYEKDVNGDEKDSSDDEDYEEE-----DEELGDMP
KSADPMEWFVYLQAQHFTLPDKVNDFGQKE---------------------------------CKKIQNARPGTIGK
YLSTIG-------------------------------------------------------------------------------------------------------------------
--------------------------------------------------------------------------------------
>Obodhiang/1-429
------------------------MFCTINQKAIRPAKPSDSTTPQYPSEFFDKNNYQRPTVRVTQGGYKIQE
LREILSN--GIL-QDDINPHHVVRYMELIMEGITDTLDDDWTSFGVKIGKKGEKITPLSLLNIMIE
EDDLIDGKRNNDVTKKD------DKWIMLIITSYYRFAFSQNPNHRSNLITKLNLQLRTFLKDPPTI
VDN--------MGLFASLISNVNFNKLISALDMFLNRFKNNDWSFLRFGTIASRYKDCSALMSLSH
VCEVTGMKIEEFMDWIFVYSTGEDMIKLMREGNEIDDPLSYMPYTMSMGLSMKSPYSSINC
PSIYSFIHMLGSLLGSERS-------------------------------------------------------RNARMVSENNIVNLKM
NAGVVSYVKSHRASMIKAFISNEVKDQWYDNEGNEADDKTDDESDDE--------LDGMPKGD
NPVEWFMFLESRHFELPDEIKSFMNRE---------------------------------AKKITNPRSGTIGKFVSL
MN-------------------------------------------------------------------------------------------------------------------------
--------------------------------------------------------------------------------
>AdelaideRiver/1-429
------------------------MFCTINQKAVQPAKPSDTTTPQYPADFFNKNNHQKPTVRVTQRGYKIQE
LREIISN--GIV-QDDLNSHHVVRYMELIMEDITDTLDEDWNSFGVKIGRKGDKITPLSLVNVMI
EEDELIDGKRNNGVNKKD------DKWIMLVITSYYRFAFSQNQNHRSNLITKLNLQLRTFLKDP
PTIVDN--------MGLFTSLISNINFTKLISALDMFLNRFKNNDWSYLRFGTIASRYKDCSALMSL
SHVCDVTGMKMEEFMDWIFVYSTGEDMIKLMKEGNEIDNPMSYMPYTMSMGLSTKSPYSS
INCPSIYSFIHMLGSFLGSERS-------------------------------------------------------RNARMVSENNIVN
LKVNAGVVSYVKSHRASMIKAFISNDVKEQWYNNDDNDNENGGDDESDEE--------LDEMPK
GDNPVEWFMYLESRHFELPEEIKNFMNRE---------------------------------ARKITNPRVGTIGKFV
STMN---------------------------------------------------------------------------------------------------------------------
------------------------------------------------------------------------------------
>Kotonkan/1-426
------------------------MFCTITETSVKAIKPTDNVPPQYPGDYFGRSKGTKPTIRIPQSKLDL

QAARELVKG--GLS-KGELSVKHGIRYLYLLMCEVNETMDGDWESFGVVIGKKGENCNPLSM
FNIIEEDDKLIDGTKNPKATPED------DKWMALAVISMYRLGRTTNQTHRNTVITKLNAQMQGI
SKDAIPMIDL--------PSLQSSWVANQDFCKIVAGIDMFFNKHKMNDWAYLRFGSIPSRFKDCS
ALLSLGHICDVTGMDLTEFLDWIFVGTVAKEVVGMMKEGNEIDNAYSYTPYMMDMGLSLKS
PYSSTVCPGTYTLVHMIGTLLFSDRS----------------------------------------------------KHAKMISEN
NLSNIRINSEVVAYVKGKKGSLVKAFIRPEFKDQYKDDDTTDSESEEGDEAGSL------------PR
TDDPMEWFAYLEYNHFDLPDVIKEHTRSE--------------------------------SRKIQNTRAGTIGNHV
VTTFN-------------------------------------------------------------------------------------------------------------
-------------------------------------------------------------------------------
>Koolpinyah/1-426
------------------------MFCTITETSIRAIKPTDNVPPQYPGDYFVRSKGSKPTIRIPQSKLDLQAAR
ELVKG--GLS-KGELSVKHGIRYLYLLMCDINEVMDEEWESFGVVIGKRGENCNPLSMYNIIEE
DDKLIDGTKNPKATADD------DKWMALAVTAMYRLGRTMNQTHRNTVITKLNAQMQGISKDA
IPMIDL--------PSLQASWVSNQDFCKIIAGIDMFFNKHKMNDWAYLRFGSIPSRFKDCSALLSL
GHICDVTGMDLTDFLDWIFVGTVAKEIVGMMKEGNEIDNAYSYTPYMMDMGLSLKSPYSST
VCPGTYTLVHMIGTLLFSDRS----------------------------------------------------KHAKMISENNLSNI
RINSEIIAYVKGKKGSLVKAFIKPEFKDHYKDDDTTDSESDDGDEAGSL------------PKSDDPM
EWFAFLEYNHFDLPDVIKEHTRLE--------------------------------SRKIQNTRAGTIGNHVVTTFN--
-----------------------------------------------------------------------------------------------------------------------
-------------------------------------------------------------------
>NewKentCounty/1-426
------------------------MFCTVTETSIKAYKPSDNVPPQYPKDYFEKNRGNKPTIRIPQSRLDL
TAARELVKG--GLS-KGDLSVKHAMRYLYLILDQVSETADSDWSSFGIRICNKNDEAKVFMMY
NVMEXBDSVPDGKXXPPSTEDD------DKWMALALVSIYRLGRTNNQAHRNLILTKLNAQLQG
INKDAVAMVDL--------PALQAAWLSNPEFCKLIAGIDMFFNKYKNNPWAFLRFGTIPSRFKDC
SALLSLGHICDITGMDLENFFEWIFVGSVAKEIVSMMKEGNEIDDAYSYMPYMMDLGLSLKS
PYSSTACPGTYTLVHTIGALLFSERS----------------------------------------------------KNARMISEN
NLSNIKINAEIVAYVKGKKGSLVKAFVKNDQKDYYKEENNSDNESEEGDEAGEL------------PS
SDDAMEWFAYLEXXHFDLPDIIKEHTRIE--------------------------------SKKLTGTRAGTIGYHIVN
TFN----------------------------------------------------------------------------------------------------------------
---------------------------------------------------------------------------
>Porcine2/1-424
------------------------MYCTVTDSVIRPKRPHDNVPAQFPKDYFSRNNHTKPTIRVPQKDLSI
QDARELVRG--GLV-RNDLNVKHAMRYMYLILAKINETAEEDWESFGIKIAQKGCQVTPWDAF
NIIEDKDKLVDGVKDNKAVDED------DKWMALSIVTTYRLARTNNQTHRNNLIVKANQQIAGM
NKDAPSLIDL--------PAQQAAWISNPDFTKMMAAIDMFFNRYKNSEWAFLRFGTIPARYKDCA
GLMSIGHLCDVTGLELDDVLDWIFVGTVATEIVNMMKEGNEVDDPYSYMPYMMELGISLKS
PYSSSMCPGLYTFVHIVGCLLYSERS----------------------------------------------------KNARMVSD
NNLVNIKMNAEVLAYVRSKKGDLVKAFVKENEKDRLEKT--TDESEIAEVDMSQM------------PT
SNDPMEWFAYLEFMDFNLPEEIKHHTKSE--------------------------------SKKITNTRAGTIGNHV
STTYC-------------------------------------------------------------------------------------------------------------
-------------------------------------------------------------------------------
>Porcine1/1-424
------------------------MYCTVTDSTIKPKRPYDNVPAQFPLDYFKRNNHTKPNLRIPQKELGL
QDVRELVRG--GLN-RNDLNVKHAMRYLYLVLSKVVETADDDWDSFGIKLVRKGCQMTPWDL
FNIVEDKDKLVDGIKDNKAVEDD------DKWMCLAIVATYRLGRTNNQAHKNNLIVKINQQLVG
MNRDAPPMIDL--------PAQQAAWVANADFNKMIAGIDMFFNKFKTNEWAFLRFGTIPARYKD
CAALMSIGHLCDVTGLDLEDVLDWIFVGTIATEIVNMMTEGNEVDNPYSYMPYMMELGLSM

KSPYSSSMAPGLYTLVHMVGCLLYSDRS-------------------------------------------------KNARMI
SDRNLVNIKMNAEVLAFVRAKKGDLVKAFVKENEKNRLEKD--SDNIEPTELDLTSL------------
PSSHDPMDWFAYLESVDFNLPDEIKLHTKSE------------------------------SRKITNTRAGTIGHH
IATTFI-------------------------------------------------------------------------------------------------------
-------------------------------------------------------------------------

>Yata/1-426
-----------------------MFSALSGKPVAACMPHETIPPQYPADFFRNNKNTKPHIRIPMSELDLDR
VRKFVHA--GIIDRNKLKLPHAMRYLFLVLSKIKEKLDNDWKSFGVTIGLKDQDLDPFCMYVV
HHDAGPQVDGTESTTVGPED------DDWMVMYLLGIYRLGRTQNDTHKGNLATRLLAQMKSL
NPRTLNITND--------DALQNIWVGNTDYCKLIAGVDMYFNRFKKSDFAYLRFGTIPSRFKDCA
SLLSIGHVCNLTGMTLEEYLGWIFVATVGREIESMMKEGNEVDQPFSYMPYMMEMGLSMK
SPYSSAASPGVYTLAHIIGTLLFSERS-------------------------------------------------KNARMVSE
NNLSNIRINAEIVAYVRARKGSLMKVFHKDKASLEAAQAVQEDNGS-VDVLLGDL------------PA
GNDPDEWFTTLEINQFELPHEIRTHTISE------------------------------SRKIGNTRAHTIGHHVAT
TFV---------
---------------------------------------------------------------------------------------------------------------
---------------------------------------------------------

>VSV/1-421
--------------------SVTVKRIIDNTVIVPKLPANEDPVEYPADYFRKSKEIPLYINTTKS---LSDLRG
YVYQ--GLK-SGNVSIIHVNSYLYGALKDIRGKLDKDWSSFGINIGKAGDTIGIFDLVSLKALDG
VLPDGVSDASRTSAD------DKWLPLYLLGLYRVGRTQMPEYRKKLMDGLTNQCKMINEQFE
PLVPEG-------RDIFDVWGNDSNYTKIVAAVDMFFHMFKKHECASFRYGTIVSRFKDCAALAT
FGHLCKITGMSTEDVTTWILNREVADEMVQMMLPGQEIDKADSYMPYLIDFGLSSKSPYSS
VKNPAFHFWGQLTALLLRSTRA-------------------------------------------------RNARQPDDIEYT
SLTTAGLLYAYAVGSSADLAQQFCVGDNKYTPDDSTGGLTTNAP-----------------PQGRDVVE
WLGWFEDQNRKPTPDMMQYAKRA------------------------------VMSLQGLREKTIGKYAKSEF
DK--------
---------------------------------------------------------------------------------------------------------------
---------------------------------------------------------

>RABV/1-450
--------------------MDADKIVFKVNNQVVSLKPEIIVDQYEYKYPAIKDLKKPCITLGKAPDLN
KAYKSVLSC--MSA--AKLDPDDVCSYLAAAMQFFEGTCPEDWTSYGIVIARKGDKITPGSLVE
IKRTDVEGNWALTGGMELTRDPTVP-EHASLVGLLLSLYRLSKISGQSTG-NYKTNIADRIEQIF
ETAPFVKIVEHHTLMTTHKMCANWSTIPNFRFLAGTYDMFFSRIEH-LYSAIRVGTVVTAYED
CSGLVSFTGFIKQINLTAREAILYFFHKNFEEEIRRMFEPGQETAVPHSYFIHFRSLGLSGKSP
YSSNAVGHVFNLIHFVGCYMGQVRS-------------------------------------------------LNATVIAAC
APHEMSVLGGYLGEEFFGKGTFERRFFRDEKELQEYEAAELTKTDVALADDGTVN---SDDE
DYFSGETRSPEAVYTRIIMNGGRLKRSHIRRYVSV------------------------------SSNHQARPNS
FAEFLNKTYSSDS-------
---------------------------------------------------------------------------------------------------------------
---------------------------------------------------------

>EBOV/1-739
-----------------MDSRPQKIWMAPSLTESDMDYHKILTAGLSVQQGIVRQRVIPVYQVNNLEEIC
QLIIQAFEA--GVDFQESADSFLLMLCLHHAYQGDYKLFLESGAVKYLEGHGFRFEVKKRDGV
KRLEELLPAVSSGKNIKRTLAAMPEEETTEANAGQFLSFASLFLPKLVVGEKACLEKVQRQIQ
VHAEQGLIQYPTAWQSVGHMMVIFRLMRTNFLIKFLLIHQGMHMVAGHDANDAVISNSVAQA
RFSGLLIVKTVLDHILQKTERGVRLHPLARTAKVKNEVNSFKAALSSLAKHGEYAPFARLLNL
SGVNNLEHGLFPQLSAIALGVATAHGSTLAGVNVGEQYQQLREAATEAEKQLQQYAESREL

DHLGLDDQEKKILMNFHQKKNEISFQQTNAMVTLRKERLAKLTEAITAASLPKTSGHYDDDD
DIPFPGPINDDDNPGHQDDDPTDSQD--TTIPDVVVDPDDGSYGEYQSYSENGMNAPDDLV
LFDLDEDDEDTKPVPNRSTKGGQQKNSQKGQHIEGRQTQSRPIQNVPGPHRTIHHASAPLT
DNDRRNEPSGSTSPRMLTPINEEADPLDDADDETSSLPPLESDDEEQDRDGTSNRTPTVAP
PAPVYRDHSEKKELPQDEQQDQDHTQEARNQDSDNTQSEHSFEEMYRHILRSQGPFDAVL
YYHMMKDEPVVFSTSDGKEYTYPDSLEEEYPPWLTEKEAMNEENRFVTLDGQQFYWPVM
NHKNKFMAILQHHQ
>Measles/1-524
ATLLRSLALFKRNKDKPPITSGSGGAIRGIKHIIIVPIPGDSSITTRSRLLDRLVRLIGNPDVSGP
KLTGALIGILSLFVE--SPG-QLIQRITDDPDVSIRLLEVVQSDQSQSGLTFASRGTNMEDEADQ
YFSHDDPSSSDQSRSGWFENKEISDIEVQD--PEGFNMILGTILAQIWVLLAKAVTAPDTAAD
SELRRWIKYTQQRRVVG-EFRLERKWLDVVRNRIAEDLSLRRFMVALILDIKRTPGNKPRIAE
MICDIDTYIVEAGLASFILTIKFGIETMYPALGLHEFAGELSTLESLMNLYQQMGETAPYMVILE
NSIQNKFSAGSYPLLWSYAMGVGVELENSMGG-------------------------------------------------------------LN
FGRSYFDPAYFRLGQEMVRRSAGKVSSTLASELGITAEDARLVSEIAMHTTEDRISRAVGPR
Q---AQVSFLHGDQSENELPGLGGKEDRRVKQGRGEARESYRETG------------------------------------
SSRASDARAAHPPTSMPLDIDTASESGQDPQDSRRSADALLRLQAMAGILEEQGSDTDTPR
VYNDRDLLD--------------------------------------------------------------------------------------------------------------------------
------------------------------------------------
>Nipah/1-531
SDIFEEAASFRSYQSKLGRDGRASAATATLTTKIRIFVPATNSPELRWELTLFALDVIRSPSAA
ESMKVGAAFTLISMYSERPGALIRSLLNDPDIEAVIIDVGSMVNGIPVMERRGDKAQEEMEG
LMRILKTARDSSKGKTPFVDSRAYGLRITDMS-------TLVSAVITIEAQIWILIAKAVTAPDTAEES
ETRRWAKYVQQKRVNP-FFALTQQWLTEMRNLLSQSLSVRKFMVEILIEVKKGGSAKGRAV
EIISDIGNYVEETGMAGFFATIRFGLETRYPALALNEFQSDLNTIKSLMLLYREIGPRAPYMVLL
EESIQTKFAPGGYPLLWSFAMGVATTIDRSMG-------------------------------------------------------------ALNI
NRGYLEPMYFRLGQKSARHHAGGIDQNMANRLGLSSDQVAELAAAVQETSAGRQESNVQ
AREAKFAAGGVLIGGSDQDIDEGEEPIEQSGRQSVTFKREMSISSLAN---------------------------------
SVPSSSVSTSGGTRLTNSLLNLRSRLAAKAAKEAASSNATDDPAISNRTQGESEKKNNQDLK
PAQNDLDFVRADV----------------------------------------------------------------------------------------------------------------
-------------------------------------------------------
>RHSV/1-390
------------------------------ALSKVKLNDTLNKDQLLSSSKYTIQRSTGDSIDTPNYDVQKHINKL
CGMLLITEDANHKFTGLIGMLYAMSRLGREDTIKILRDAGYHVKANGVDVTTHRQDINGKEM
KFEVLTLASLTTEIQINIEIESR-------KSYKKMLKEMGEVAPEYRHDSPDCGMIILCIAALVITKL
AAGDRSG-----------LTAVIRRANNVLKNEMKRYKGLLPKDIANSFYEVFEKHPHFIDVFVHFGI
AQSSTRGGSRVEGIFAGLFMNAYGAGQVMLRWG-----VLAKSVKNIMLGHASVQAEMEQVV
EVYEYAQKLGGEAGFYHILN-----------------------------------------------------------------NPKASLLSLTQFPH
FSSVVLGNAAGLGIMGEYRGTPRNQDLYDAAKAYAEQLKENGVINYS------------VLDLTAEE
LEAIKHQLNPKDNDVEL---------------------------------------------------------------------------------------------------------
-----------------------------------------------------------------------------------------------------------------------------------------------
------------------------------------------------
